# Dietary road salt and monarch butterflies: minimal effects on larval growth, immunity, wing coloration, and migration to Mexico

**DOI:** 10.1101/2023.09.04.554310

**Authors:** Amanda K Hund, Timothy S. Mitchell, Isabel Ramirez, Amod Zambre, Lili Hagg, Anne Stene, Karilyn Porter, Adrian Carper, Lauren Agnew, Alex Shephard, Megan Kobiela, Karen Oberhauser, Orley R. Taylor, Emilie Snell-Rood

## Abstract

The spectacular migration of the monarch butterfly is under threat from the loss of habitat and the decline of their milkweed host plants. In the northern part of their range, roadsides could potentially produce millions of monarchs annually due to high densities of milkweed, however roadside milkweed can accumulate chemicals from roads, such as sodium from road salt. Controlled lab studies have shown mixed effects of sodium on monarch development: small increases can be beneficial as sodium is an important micronutrient in brain and muscle development, but large increases can sometimes decrease survival. It is unclear how dietary sodium affects performance in ecologically relevant conditions, and the migration itself. In this experiment, we raised monarchs outdoors, in migration-inducing conditions, on milkweed sprayed with three levels of sodium chloride. We released 2500 tagged monarchs and held an additional 250 for further lab assays. While our recovery rates to the wintering grounds were low (N = 7 individuals), individuals from all three sodium chloride treatments made it to Mexico. Butterflies reared on control milkweed and low salt concentrated sodium in their tissues, while those on high salt diets excreted sodium, suggesting levels were above a physiological optimum. There were no effects of treatment on wing coloration, survival, body size, immunity, or parasite prevalence. Taken together, our results suggest that monarchs are robust to levels of sodium in milkweeds found along roadsides, which is promising with respect to monarch conservation efforts that promote roadside habitat.

**Significance Statement:** Monarch butterflies are a flagship species for pollinator conservation, and were recently being listed as endangered by the IUCN. Roadside habitat is a target for monarch breeding habitat as they often have high densities of milkweed, the monarch hostplant. However, roadsides can also have high levels of pollutants, such as salt from deicing treatments. We reared monarch caterpillars on sodium treated milkweeds, measuring a suite of performance measures, and releasing nearly 2500 tagged monarchs for migration. We found little effect of salt on migration to Mexico, survival, body size, development time, parasite prevalence, immunity, or coloration. Monarchs appear robust to levels of sodium found in milkweed along roadsides, supporting the possibility of roadsides as habitat.

## Introduction

The monarch butterfly, *Danaus plexippus*, (Lepidoptera: Nymphalidae) is one of the world’s most charismatic insects and acts as a flagship species for conservation [1–3]. One of the most fascinating aspects of the monarch’s life cycle is the annual migration, where, in the eastern and central regions of North America, individuals fly up to 4000-km at the end of each summer –– from the northern United States to over-wintering grounds in Mexico [4–6]. In the spring, monarchs move back into their breeding grounds, following the northward phenology of their milkweed host plants, *Asclepias*, spp. (Apocynaceae), over 3-4 consecutive generations.

However, monarch populations are under threat due to habitat loss, pesticide use, climate change, and the spread of disease [7–9]. In many ways, they are a prominent example of concerns of a broader “insect apocalypse” [10–12]. The decline of monarchs has led to their wait-listing as threatened by the USFWS and their global listing as endangered on the IUCN list [13, 14]. Many initiatives seek to aid monarch conservation by increasing host plant availability and leveraging pockets of potential habitat [15–18], though evidence-based management has yet to keep up with all initiatives.

One of the most controversial monarch conservation initiatives has been the push to restore roadside habitat for monarchs. Roadsides represent a promising opportunity for monarchs, as they potentially capitalize on millions of acres of underutilized habitat and can act as corridors for insect movement [19–22]. Recent legislation has set aside funds for roadside restoration for monarchs [23], and some states prioritize roadside restoration as green infrastructure (e.g., Iowa Living Roadway Trust Fund). Roadsides in the Midwestern United States often contain thousands of milkweeds along every kilometer of roadside [24, 25], and can support high densities of developing monarch larvae [26, 27]. However, roads come with substantial risks, including vehicle collisions [28–30] and pollutant exposure [31–33].

Sodium chloride is a pollutant of concern in many parts of the monarch’s breeding range due to the application of road salt as a winter deicing agent. Sodium, as a positively charged cation, tends to associate with negatively charged soil particles, accumulating in areas with high salt application [34, 35], while chloride leaches more rapidly into waterways where it takes a negative toll on aquatic life and accumulates as a legacy pollutant [36–38]. Sodium can be taken up by roadside plants or adhere to leaves as dust [33, 39]; in either case, it can be consumed by developing insect larvae on roadside plants [33]. Sodium is an important micronutrient in animal development, playing a key role in neural and muscle development and function [40–42].

However, high amounts of sodium can be toxic to animals, including mammals [43–47] and insects [48–52]. Laboratory rearing studies have reflected this dual role of sodium as both a nutrient and toxin, with mixed results on monarch performance, sometimes finding beneficial effects on eye and muscle development [42], sometimes highlighting concerns for survival [53], and sometimes finding no effect [54]. Given that many toxicity studies are confined to laboratory conditions, it is hard to know how these diverse effects of sodium play out in more field-realistic conditions, and in the monarch migration itself. Elevated sodium may improve migratory performance if it affects navigation or flight abilities through impacts on sensory systems or flight muscle [42]. However, high levels of sodium could result in physiological stress that could lengthen development time, or decrease flight performance and survival [53].

To test the net effects of sodium chloride on monarch performance in the field, we took advantage of the Monarch Watch Program, an initiative started in 1992 that relies on a network of scientists and monarch enthusiasts that have tagged over half a million migrating monarchs with small wing stickers [55]. The recovery rate of midwestern monarch tags at the wintering grounds in Mexico averages 1.4%, although migratory performance and recovery rates for cage-reared monarchs are lower [56, 57]. We aimed to rear 3000 monarchs under three sodium chloride treatments, and test differences in recovery at sites 3000-km away in Mexico (El Rosario and Cerro Pelon). We measured body sodium content of our experimental butterflies relative to their diet, and to individuals from the wintering sites, as a measure of sodium requirements in development (the “bioconcentration factor” [58]). At the same time, we assayed a wide range of performance metrics on our reared butterflies to determine effects of salt exposure in field-realistic conditions. Specifically, we measured development time, larval growth rate, and adult body size, as timely emergence and body size are associated with migration and recovery at the wintering grounds [59–62]. We took measures of wing coloration, as darker, redder wings are associated with migratory status and flight performance [63, 64]. We also assessed infections of the parasite *Ophryocystis elektroscirrha* (“Oe”) [65], which has negative effects on monarch body size, survival [66], flight performance, and wing strength [67, 68]. We measured survival in climate chambers with both constant and variable temperatures. Finally, we took four measures of immune performance (hemocyte counts, filament melanization, phenoloxidase activity, and prophenoloxidase activity [69]) given past data suggesting relationships with immune capacity and migration [70].

We varied sodium exposure across three levels, meant to simulate variation in sodium exposure across Midwestern roadside milkweeds, where sodium levels are elevated in plants next to the road edge or on high traffic roads [33, 71]. Our low sodium (360 ppm) and high sodium (2171 ppm) treatments approximated the 90th and 99th percentiles of milkweeds collected from across Minnesota roadsides, respectively (246 and 2300 ppm [33, 71]). Our control treatment (113 ppm Na) made use of milkweeds collected from >50m from the roadside, treated with distilled water only. We predicted that monarchs in the low sodium treatment would show the highest performance in field-realistic conditions, and during the migration itself, as they would have access to a limited micronutrient, but below stressful levels.

## Methods

### Overview and rationale

In the eastern US population of monarchs, migratory behavior and reproductive diapause is induced by a suite of cues at the larval stage, including declining photoperiod, low temperatures at night, and to some extent low host plant quality [72–74]. Given this range of factors, inducing migratory status in climate chambers or greenhouse conditions is challenging (see variation in [73]), and the most reliable way is to rear outdoors, in natural conditions. Additionally, outdoor rearing simulates natural conditions as much as possible, including fluctuations in temperature and humidity, allowing us to test how previous lab rearing under different sodium chloride treatments [42, 75] may translate to semi-realistic field conditions. To rear monarchs outdoors, we chose a site on the University of Minnesota-St Paul campus, nested within experimental agricultural fields, but surrounded by trees to provide shaded buffering of daily heat extremes in the rearing cages (exact coordinates: 44.988241, –93.177998). We sought to time our release with the leading edge of the monarch migration as recovery rates tend to be higher for this first wave of migrating butterflies [56, 76]. Based on average weather conditions and the timing of the declining sun angle, this tends to be around August 20-25th in our region, so we planned the timing of our rearing accordingly (Figure S1). Our peak release dates were about a week later than projected (Figure S2) due to lower than normal August temperatures.

### Larval rearing and dietary treatments

Full details of rearing can be found in the supplemental methods. We first raised a generation of “breeder monarchs” (F1) from wild-caught larvae and eggs. In total, 140 of these common-garden reared individuals emerged as adults (from July 26 2019 to July 28 2019) and were used to found the experimental generation. To rear experimental butterflies (F2), we collected eggs (>5000) from over 50 common-garden-reared females over a two-week period (July 19-July 31). Each day, greenhouse-collected eggs were counted and moved to 0.1 m^2^ cages held at room temperature (approximately 23C) in a windowsill for 5-6 days before larvae were transferred to outdoor cages. Larvae from a given egg collection cage were divided between the three sodium chloride treatments in outdoor cages (61×61×61cm) staked to the ground. Milkweed supply in outdoor rearing cages was refreshed daily (1-10 stalks each day, depending on age of larvae) with common milkweed (*Asclepias syriaca*) collected from Cedar Creek Experimental Research Station (approximate coordinates: 45.405401, –93.186604), which generally has milkweeds with low sodium concentrations (based on [42, 75]).

Experimental larvae (F2) were reared on one of three diet treatments starting at their day of transfer to outdoor conditions (Figure S1, S3). The control treatment was milkweed sprayed with distilled water. The low treatment was milkweed sprayed with a 0.24g NaCl per 25-fl-oz (739 mL) solution, meant to mimic somewhat elevated sodium in roadside plants [33, 71]. The high treatment was milkweed sprayed with 2.1 g NaCl per 25-fl-oz (739 mL) solution, meant to mimic high sodium plants from close to the road edge [33, 71]. We confirmed the target sodium concentration of our manipulation by randomly selecting three treated milkweeds from each treatment to measure sodium concentration using ICP-AES ([77] at UMN’s Research Analytical Lab, RAL). Milkweeds from each treatment group showed significant variation in sodium concentration (F_2,6_ = 77.5, P < 0.001), with those in the high group having significantly higher sodium (2171 ppm) than the control (113 ppm Na) and low (360 ppm Na) treatments. These levels approximate the average, 90th and 99th percentile of milkweeds collected along Minnesota roadsides [33, 71] and were in line with our target sodium concentrations in the design of the experiment (50-500-2000 ppm). We also measured nitrogen content using the Dumas Method ([78] at UMN-RAL) and saw no significant variation across treatment groups (F_2,6_ = 1.93, P = 0.22), with an average nitrogen content of the milkweeds at 3.06%.

### Emergence, tagging, release, tag recovery

Within 24 hours of emergence, adult monarchs were marked with a fine-tipped black sharpie with an individual number, sexed, imaged, and screened for Oe (see below). An individual’s de development time was defined as the number of days between egg collection and adul emergence. After processing, adult butterflies were held in a large screened tent (Coleman screen house: 3m x 3m x 2.1m) with ad lib access to a sugar solution (per 2L distilled water: 300g sucrose, 8g ascorbic acid, 4g sorbic acid, 4g methylparaben, 3 pinches pollen) provided in a dish with colorful nylon scouring pads. The following day, each individual butterfly received a Monarch-Watch tag on the discal cell of the ventral hindwing, according to tagging protocols [55, 76] and were released (generally between 8:30 and 11 am). Any butterflies that emerged after September 11, 2019 were assumed to be too late to likely reach the wintering grounds in Mexico [76]. Indeed, our recovered individuals had an average release date of Sept. 3^rd^ (latest Sept 7^th^). The 246 butterflies that emerged after September 11 were primarily from the final set of 20 rearing cages set up outside (July 27 and later, 80% of late individuals). These butterflies went into several groups to measure a range of other phenotypic traits that might differ with sodium chloride treatment, including immune measures and thorax sodium levels. Tags were recovered by the Monarch Watch program during visits to wintering grounds in 2020 and 2021; tags are generally found on dead individuals or on the ground.

### Survival in lab

To measure survival in the lab, butterflies (N = 152) were held in mesh cages (30 x 30 x 30 cm; N < 20 individuals per cages) in low light conditions in one of two climate chambers. Adult butterflies were sprayed daily with distilled water but did not have access to food. Butterflies were assigned randomly to one of two temperature treatments (constant 21C, or variable with 21C daytime and 7C nighttime temperatures) to measure consistency in survival differences across different conditions. Butterflies were placed into the chamber on the day of their emergence and cages were checked daily for deaths. Temperature treatments were chosen to mimic average September outdoor conditions in Minnesota, with one chamber set around the average high (at constant conditions) and one chamber set at the average high with a decrease at night to average September lows (7C). Previous work had found that cool temperature swings might bias survival across sodium chloride treatment groups [75].

### Thoracic sodium content

We used the sodium content of an adult butterfly’s thorax, relative to the sodium content of their diet, as a measure of whether sodium was limited in availability, or at stressful levels. We chose to focus on the thorax as a representative tissue as it is primarily flight muscle, which would be heavily influenced by larval diet, and also be functionally reliant on sodium for muscle function [42]. Individuals from the three rearing groups were sacrificed at –20C on the day of emergence. Thoraxes were removed from the body and wings and dried at 70C for 48 hours. Thoraxes were processed for sodium content at the Quantitative Bio-element Imaging Center (QBIC) facility at Northwestern University using ICP-MS with a digestion in nitric acid (250 ul) and hydrogen peroxide (63 ul; see supplementary materials for more details). Sodium content was calculated as a function of thoracic weight, as parts per million.

### Collection of monarchs on the wintering grounds

To compare the thoracic sodium content of reared monarchs to those individuals that made it to Mexico, one team member (I.R.) collected 40 wild individuals (not from our experiment) from Cerro Pelon on January 21, 2021. The thorax was separated from the body and initially dried in the field in a desiccation chamber. Samples were processed at the QBIC facility (see above) with ICP-MS, as for reared samples. We preferred non-destructive sampling of dead individuals, but to ensure we did not have a bias due to selection of dead individuals, we sampled an even mix of dead and alive individuals. There was no significant effect of “dead” or “alive” on thoracic sodium content (in a model controlling for sex, F(1,36) = 0.0006, P = 0.98), so we merged all individuals for this analysis.

### Weather Data

Because temperature is important in the development of insects, and humidity could influence how dietary sodium is expelled, we included weather data in many of our statistical models. We used hourly weather data from the weather station at the University of Minnesota St. Paul Campus Climate Observatory, which was ∼480 meters from our outdoor rearing site. For each individual, we took the average of the hourly weather data during the days that individual was developing (from transfer to eclosion). To account for both temperature and humidity, we calculated vapor pressure deficit (VPD). As VPD was tightly correlated with both temperature and humidity data, we used just VPD in our statistical models (see figure S4).

### Morphological measurements

During adult processing, we took two images of each butterfly’s forewing for later morphological measurements. We used a Canon-EOS T3-Rebel DSLR with a 50-mm macro lens, mounted on a tripod. The butterfly was placed in a standardized lightbox (470 Studio) with two LED light strips (5500K) plugged into a portable, external power-bank. An “X-rite classic color checker” was included in each image to standardize light conditions for color measurements. For the body size measurement, we imaged the ventral side of the forewing, including the wing apex and articulation with the thorax in the photo, holding the butterfly flat for the image. We used these digital images to measure wing length using Image J (NIH). We used the standard wing length measurement used by Monarch Watch in order to calibrate our measurements with other studies [79]. For the wing coloration measurement, we imaged the dorsal side of the forewing, holding the butterfly with wings mostly closed, but the discal cell protruding for the lower wing (see figure S5).

### Wing Coloration

We analyzed several different metrics of wing coloration: the proportion of black on the wing, which is important for wing strength and durability and has been found to increase in migratory individuals; and the intensity (saturation) and redness (hue) of the orange coloration, which has been linked to mating success in males, and is associated with longer flight distances [63, 64, 80, 81]. We quantified wing coloration for the 73 individuals that also had immune data and for 276 other individuals randomly chosen from the larger dataset. Using our wing photos, we used a standardized portion of the dorsal forewing (see figure S5) and used pavo2.0 package in R to quantify the proportion of the wing that was black and MARLAB to quantify the saturation and hue of the orange color (see supplementary materials for more detail).

### Scanning for Ophryocystis elektroscirrha

Every emerged adult monarch was assayed to detect spores of the protozoan parasite *Ophryocystis elektroscirrha* (“Oe”). Each individual was held with abdomen up and a piece of scotch tape pressed gently around the edges of the abdomen before being placed on a white index card with corresponding individual label [58]. Cards were stored for later imaging on a Leica Stereo microscope at 30-150X magnification. Each Oe scale sample was independently scored by two separate researchers who were blind to treatment and to the other score for that individual and any sample with discordant scores was rescreened.

### Immune Measures

We measured four standard immune assays: filament melanization, hemocyte counts, and phenoloxidase (PO) and prophenoloxidase (ProPO) enzyme activity for 76 butterflies (25 control, 26 low, and 25 high), two days post emergence. All individuals used for immune assays emerged between 9/6-9/9/19. We collected hemolymph for hemocyte counts and to quantify phenoloxidase activity by making a small puncture on the side of the abdomen. We then inserted a roughened 2mm knotted monofilament and butterflies were held for 1 hour at 24C to quantify the encapsulation response. We photographed filaments with a black and white color standard under 20× magnification and used Adobe photoshop to quantify the degree of melanization (following [82], figure S6). We quantified the density of hemocytes using a brightline hemocytometer under 40X magnification and the enzyme activity of PO and ProPO using an absorbance spectrometer plate reader (Tecan) at 490nm using 2mM dopamine as a substrate. We measured the amount of protein in the sample using a Bradford Coomassie Plus kit (Fisher Scientific). We then calculated the activity of PO or ProPO per min per ug of protein as a measure of immune capacity (see supplementary materials for more details on immune assays).

### Data Analysis

We performed all statistics in R version 4.3.0 [83]. We report significant results in the main text and all other results in the supplement. We compared recovery rates of tagged individuals during migration using a chi-square test. We asked whether monarchs were concentrating or excreting sodium in each treatment by comparing the ratio of thoracic sodium to dietary sodium. We looked at the effect of sex and treatment on this ratio using a linear model. We also tested whether each treatment group was significantly different from one (>1 = concentrating, <1 = excreting) using one-sided Wilcoxon tests. We compared the concentration of thoracic sodium between our treatment groups and wild individuals from Mexico using a linear model, and Tukey’s pairwise comparisons. Because we found a significant effect of sex in this model, we ran models for each sex separately to look at the effect of treatment.

To test for an impact of treatment on wing length, development time (egg to adult), and growth rate (wing length / development time), we built linear mixed-effects models with sex, treatment, and mean VPD as fixed effects and larval cage ID and emergence date (Julian) as random effects using *nlme* [84]. For our wing length models, we also included the identity of the measurer as an additional random effect. We first built models with a three-way interaction between sex, treatment, and VPD. When three-way or pairwise interactions were not significant, we simplified the model to improve interpretation. For models where sex was significant (wing length, growth rate), we ran separate models for each sex to further assess any impact of treatment.

We measured survival at the level of cages (N = 60 outdoor cages) as the proportion of transferred larvae that survived to adult emergence. As we moved entire stalks of milkweed into cages and counting of small larva was difficult, this measure had some degree of error (with proportion surviving >1 in some cages), which we assumed to be random with respect to treatment. We used a linear mixed-effects model to look at the effect of treatment on survival percentage with cage and transfer date as random effects. To compare the proportion of obviously diseased larva or pupa by treatment group, we built a general linear mixed model with a beta distribution using *glmmTMB* [85], including both cage and transfer date as random effects. To assess survival in our two indoor climate chambers we used *coxme* [86] to run mixed effects cox proportional hazard models with wing length, sex, and treatment as fixed effects and larval cage as a random effect.

To compare our measures of wing coloration across treatment groups, we built mixed effects models with cageID and emergence date as random effects (modeling proportion black as a beta distribution (*glmmTMB)*, and orange saturation & hue as linear models (*nlme*). Monarch wing coloration has been known to vary with developmental temperature [87], so we included mean temperature as a fixed effect in all color models. Because color varies by sex in monarchs (females have darker coloration and wider wing veins) we modeled males and females separately. We compared OE infections using a binomial mixed-effects model with sex and treatment as fixed effects and cage as a random effect using *lme4* [88]. For our four measures of immune capacity, we built mixed-effects models with treatment and sex as fixed effects and cage and emergence date as random effects (hemocyte counts = zero-truncated poisson model (*glmmTMB,* [85]), filament darkness = gamma model (*lme4)*, PO and ProPO = linear models (*nlme*)). For models where sex was significant (hemocyte counts), we ran separate models for each sex to further assess any impact of treatment.

## RESULTS

### Migratory Recovery

Between August 15 and September 11, 2019, we released 2,464 tagged monarchs, including 831 controls, 837 individuals reared on low salt milkweed and 796 individuals reared on high salt milkweed. All domestic recoveries of released individuals (N = 14) were within the Twin Cities metro area. There were no significant differences across treatments in local recoveries (X2 = 2.12, P = 0.35), with sightings of three controls, four individuals reared on low salt diet, and seven individuals from the high salt diet (Figure 1B). Seven tags were recovered in Mexico, a total recovery rate of 0.28%. There was no differential recovery across treatment groups (X2 = 0.25, P = 0.88), with two controls, three low salt, and two high salt-reared individuals, although our sample size is obviously limited (Figure 1C).

**Figure 1.**
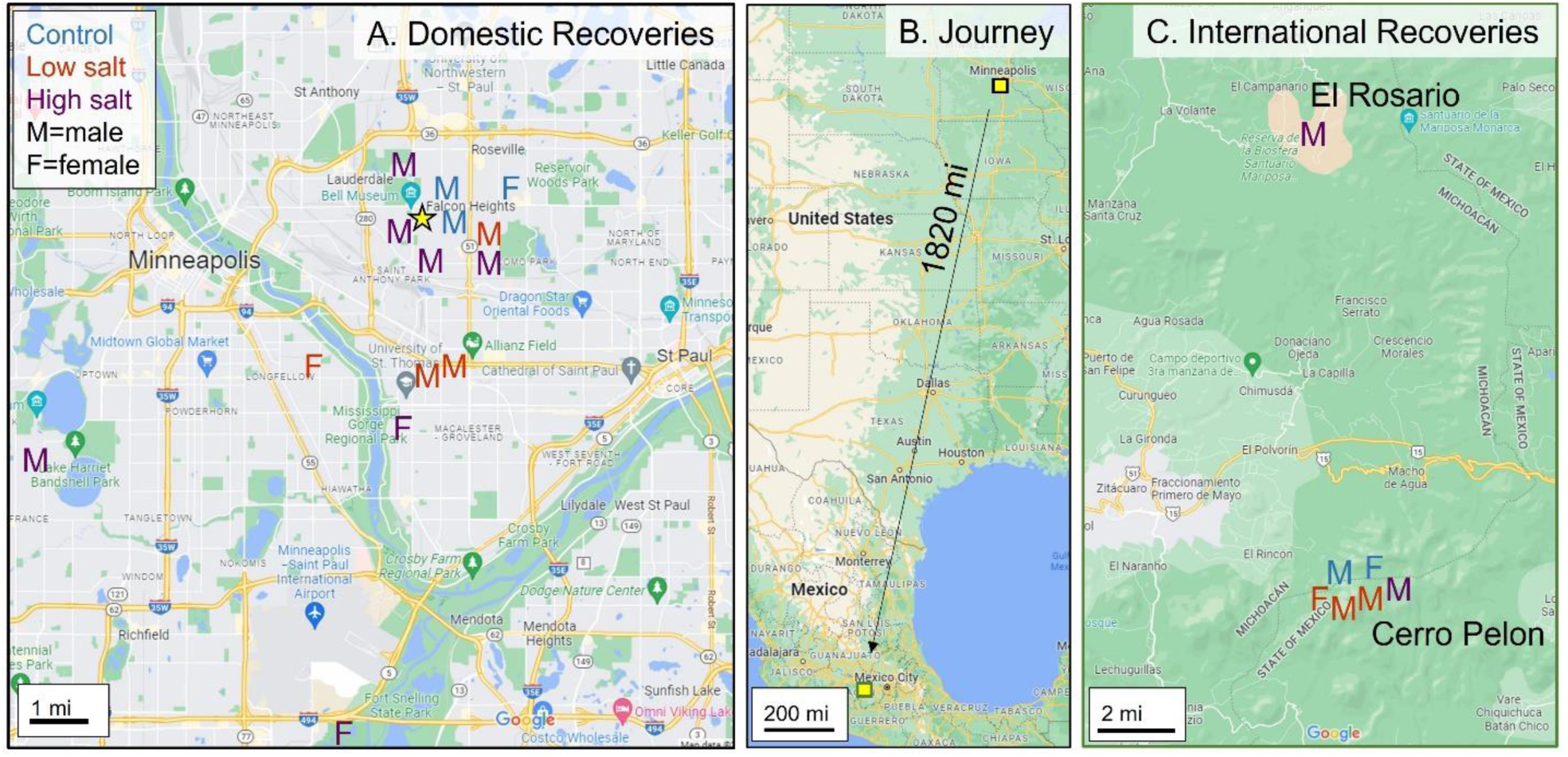
Map of recoveries. (A) Locations of domestic recoveries, all within the Twin Cities, relative to the release site (yellow star) on the University of Minnesota St Paul campus. Males are indicated with an “M” and females with an “F.” Control individuals are shown in blue; those reared on low salt diet in orange and high salt diet in purple. (B) Locations of release in the Twin Cities, MN, USA and recoveries at two wintering sites in Mexico, approximately 1820 miles away. (C) Locations of recoveries in Mexico at Cerro Pelon (6 individuals) and El Rosario (1 individual). Coding scheme corresponds to panel A.

### Sodium content of adult thoraxes

The ratio of thoracic sodium to the average sodium of each treatment diet can be used to infer whether monarchs are concentrating sodium or expelling sodium, indicative of sodium stress. We found a significant effect of treatment (ß_High_= –5.0810, SE= 0.21, t=-23.844, p<0.001; LSM: Control= 5.479, Low= 1.906, High= 0.398, Figure 2A), but no effect of sex (or sex-by-treatment) in the ratio of thoracic sodium to dietary sodium. When we compared each treatment to a value of 1 (above one is concentrating, below one is excreting) we found that butterflies in the control and low treatment were both concentrating sodium while butterflies in the high treatment were excreting sodium (Control: V= 55, p= 0.002; Low: V= 55, p= 0.002; High: V=0, p= 0.002, Figure 2A). Comparing thoracic sodium concentration of reared individuals to those collected in Mexico, we found a significant effect of both sex (ß_M_= 106.48 SE= 40.66, t=2.62, p=0.01; LSM: F= 626, M= 733) and treatment (ß_High_= 233.19, SE= 75.75, t=3..08, p=0.0003; LSM: Control= 619, Low= 693, High= 852, Wintering= 553, Figure 2B), no effect of the sex-by-treatment interaction. When we ran separate models for each sex, we found that for both males and females, treatment was significant in our model (*males:* ß_High_= 288.75, SE= 126.63, F= 2.95, p = 0.048; *females:* ß_High_= 177.12, SE= 93.47, F= 6.81, p = 0.001), more specifically the high treatment had significantly higher thoracic sodium concentration compared to measures from overwintering monarchs for both sexes (Figure 2CD).

**Figure 2.**
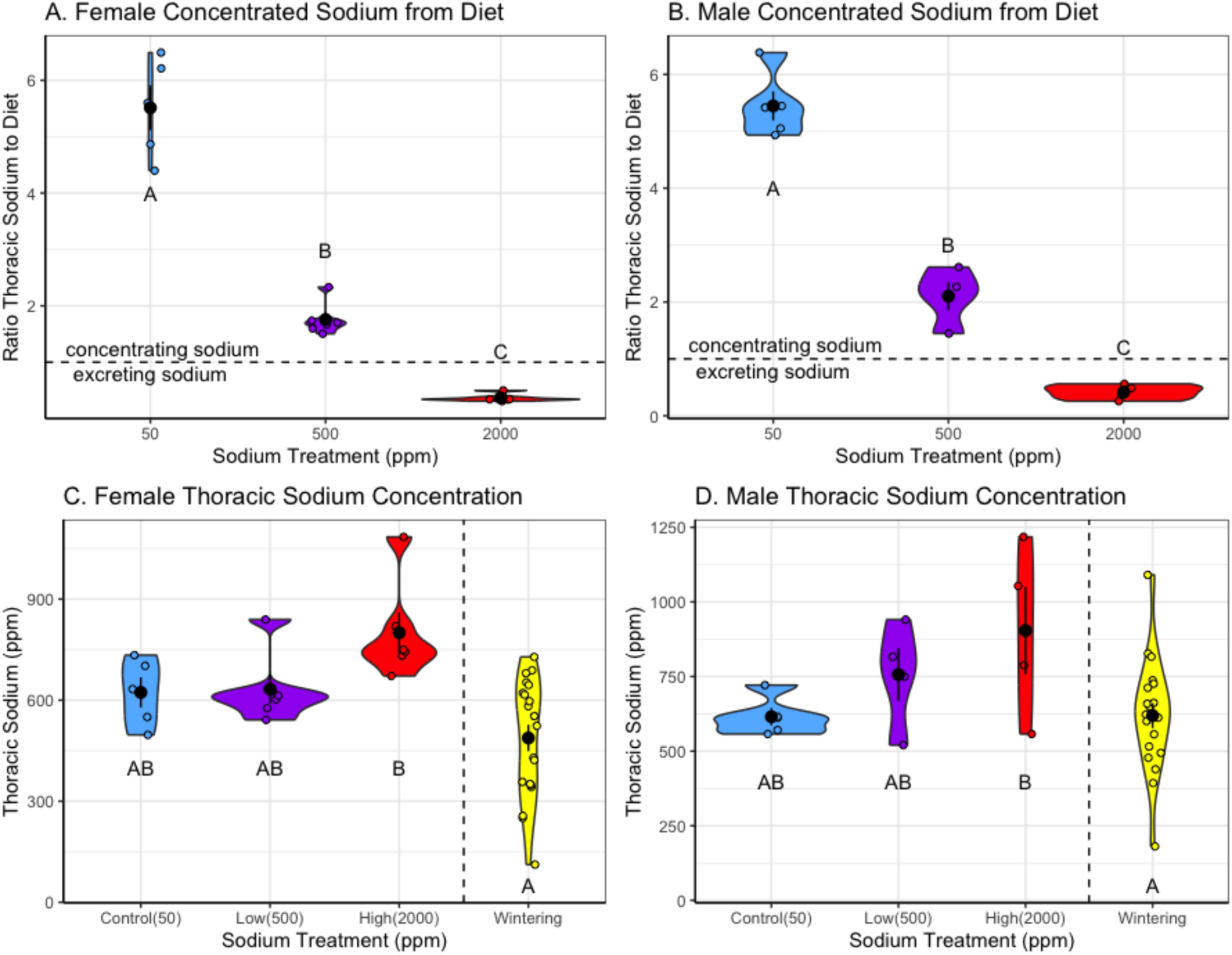
Monarch thoracic sodium content. The ratio of thoracic sodium to the average sodium concentration of each treatment diet. Values above the dotted line indicate that butterflies are concentrated sodium while values below the dotted lines indicate that monarchs are excreting sodium. Sodium chloride treatments were also significantly different from each other for this ratio for both females (A) and males (B). (C) Comparison of thoracic sodium content for females reared in experiments relative to females collected on the wintering ground. The high treatment was significantly different from wintering. (D) Comparison of thoracic sodium content for males reared in our experiments relative to males collected on the wintering ground. The high treatment was significantly different from wintering. Letters indicate significant differences using Tukey pairwise contrasts.

### Growth and development

For wing length, we found no significant interaction effects, so simplified to a model with no interactions. We found that both sex (ß= 0.64, SE= 0.092, F=48.26, p< 0.0001) and mean VPD (ß= –0.008, SE= 0.002, F=16.06, p= 0.0001) were significant predictors of wing length, where males had longer wings compared to females and wing length decreased as VPD increased (figure S7). In our separate models for males and females, we again found that VPD was significant (*females:* ß= –0.008, SE= 0.002, F=12.29, p= 0.0005; *males:* ß= –0.010, SE= 0.002, F=16.06, p= 0.0001). We found no significant effect of treatment in any of our models but did notice a trend for males where wing length was shorter in the high treatment compared to the control and low treatment (DF= 371, F=2.53, p= 0.08, control= 50.4, low= 50.7, high= 50.0) (Figure 3AB).

For development time (number of days between egg collection and adult eclosion), we also found no significant interactions. As with wing length, mean VPD was significantly correlated with development time (ß= –0.15, SE= 0.002, F=4277.44, p< 0.0001), where butterflies developed faster when VPD was greater (Figure S7). We found no effect of either treatment or sex on development time (Figure 3CD). We found similar results for growth rate, where both sex (ß= 0.015, SE= 0.002, F=41.55, p< 0.0001) and mean VPD (ß= 0.004, SE= 0.00, F=1103.77, p< 0.0001) were significant predictors of growth rate (Figure S7). Males had faster growth compared to females and growth rate increased with increasing VPD. In our separate models for each sex, we again found that VPD was significant (*females:* ß= 0.004, SE= 0.00, F=759.4, p< 0.0001; *males:* ß=0.004, SE= 0.00, F= 661.02, p< 0.0001). We found no significant effect of treatment in any of our models (for full results see tables S1-S7).

### Larval survival in outdoor cages and adult survival in climate chambers

While there was a trend of decreasing survival in the field as dietary sodium increased (*Control:* µ= 89.64%*; Low:* µ= 83.52%*; High:* µ= 78.89%), we found no effect of treatment on survival rate in our outdoor cages or on the proportion of diseased larvae and pupa found, but we acknowledge that these data were quite noisy (see tables S8-9, figure S8). For our indoor climate chamber survival study, we found no effect of treatment or sex on survival in either constant (21C) or variable (21C, 7C) conditions. However, we did find that in both conditions individuals with larger body sizes survived longer (*constant:* wing length β = –0.12, se = 0.062, Chisq = 3.67, p = 0.055; *variable:* wing length β = –0.20, se = 0.043, Chisq = 22.25, p < 0.001), and that, overall, the variable temperature conditions promoted longer lifespans (see tables S10-11, Figure S9).

### Wing Coloration

There was no effect of treatment on the proportion of black, the saturation or the hue of the orange coloration on the wing for either males or females. We did find a significant effect of temperature for orange saturation for both females (ß= –2.70, SE= 0.65, F = 17.31, p<0.001) and males (ß= –0.83, SE= 0.43, F = 3.75, p = 0.055), and for orange hue for both females (ß= – 1.45, SE= 0.42, F = 11.80, p<0.001) and males (ß= –1.73, SE= 0.33, F = 17.31, p<0.001) (see tables S12-17 and figure S10).

### Ophryocystis elektroscirrha Infections

We found no effect of treatment or sex (or treatment x sex interaction) on the number of Oe infections (*Control:* 1.81%, *Low:* 1.57%, *High*: 1.37%, see Figure S5). The overall rate of Oe detected in our experiment was 1.59% (see table S18 and figure S11).

### Immune Measures

For hemocyte counts, we found a significant sex by treatment interaction (Chisq= 326.02, p< 0.001). However, when we analyzed our sex-specific models, we found no significant effect of treatment for either males or females. For filament darkness and PO activity, there was no significant effect of sex or treatment (or a treatment x sex interaction). We found the ProPO activity was significantly different by sex (ß=-0.27, SE= 0.14, F= 3.96, p=0.05), where females had higher ProPO activity compared to males, but found no effect of treatment (or a treatment x sex interaction). When we looked at our sex specific models for ProPO we also found no significant effect of treatment (see table S18-S26 and figure S12).

## Discussion

We were interested in the net effects of sodium chloride on monarch butterfly performance in ecologically realistic conditions and the migration itself. On one hand, sodium is an essential micronutrient, and small increases can have beneficial effects on neural and muscle tissue. On the other hand, higher levels of sodium can be physiologically stressful, and high sodium diets can decrease survival. How do these joint effects play out in the field, where monarch butterflies are flying thousands of miles to reach their wintering grounds? Our results suggest that increasing salt levels in the larval diet have minimal net negative effects on monarch performance.

There was no significant variation across our treatments in survival to adulthood (Figure S6), with 2959 butterflies surviving to adulthood and 2464 tagged and released in our outdoor experiment (Figure S2), and no survival difference between treatment groups in our climate chamber study (Figure S7). In total, seven of our tagged butterflies were recovered (dead) at the wintering grounds in Mexico (Figure 1). All three treatment groups made it to the wintering grounds, including two of the high sodium chloride treatment. The recovery rate (0.3%) was less than the expected 1.4% for midwestern butterflies, consistent with lower recoveries for reared butterflies. [56, 57]. A few factors likely contributed to our realized recovery rate. First, butterflies in the experiment had an average wing length of 50.12 mm (Figure 3), slightly shorter than average wing lengths of migrating monarchs in the wild that were measured with the same methods (e.g., 51.3 and 52.1 mm, range 40-59 mm, in two Virginia populations in [59]). Reared butterflies tend to be smaller than wild monarchs, in this experiment likely due to high rearing density; this is important for recovery rates as recovery rates are higher for butterflies with longer wings [59–62]. Second, a cooler than average late August contributed to a longer development time than initially calculated for the experiment. Development time averaged 40.04 days from egg to adult, increasing over the course of the experiment, and did not differ across treatment groups (Figure 3). We had aimed to time our release with the average leading edge of the monarch migration in the Twin Cities, MN (August 20-25), but slower development times meant we were off our target by about 10 days (Figure S2), which likely also reduced recovery rates due to a delayed migration. It is likely that unusually warm conditions during the migration further reduced recovery rates for the 2019/2020 season (O.R.T.).

**Figure 3.**
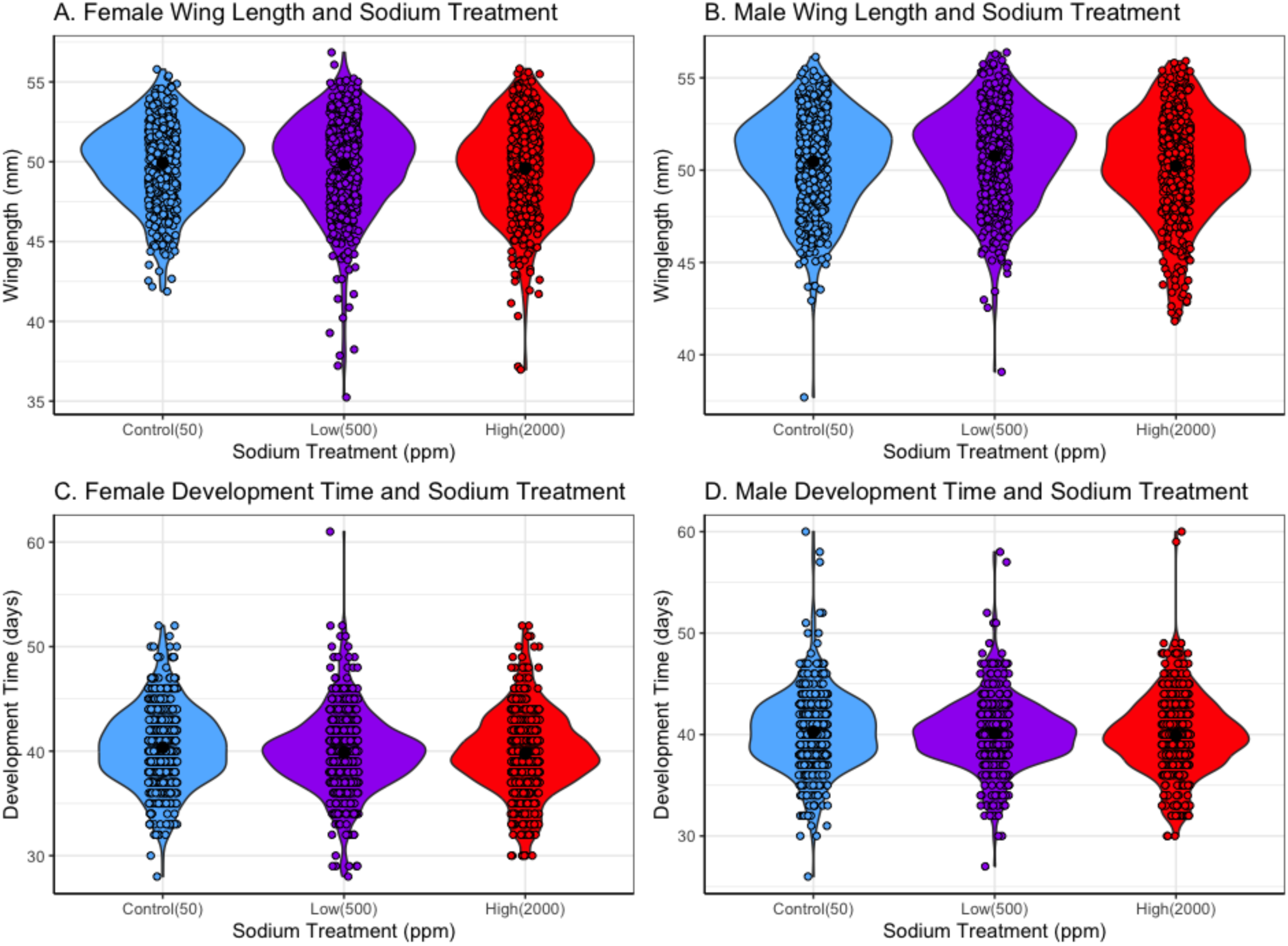
Monarch Growth and Development. There was no significant difference in wing length across sodium chloride treatments for either females (A) or males (B). Development time (days from egg collection to adult eclosion) also did not differ by treatment for females (C) or males (D).

Analysis of monarch butterfly sodium content gave additional clues to the sodium needs of migrating butterflies. For our experimentally reared butterflies, we compared the sodium content of their thorax with that of their diet, to estimate whether they were concentrating or excreting sodium. For both the control and low treatments, butterflies were concentrating sodium in their tissue relative to their diet while in the high sodium treatments, data were consistent with monarchs excreting sodium (Figure 2). In a previous experiment using similar rearing methods, monarchs similarly concentrated sodium at low sodium concentrations (75 ppm) and excreted it at high concentrations (around 2000 ppm, [54]). Insects excrete excess sodium through the actions of the Malpighian tubules, where sodium is actively pumped, and water follows [89]. Given this, we also expected sodium to be more stressful under more dry conditions (higher vapor pressure deficit), and while we saw smaller body size under such conditions, this did not vary with treatment (Supplemental Figure 7). We next compared the thoracic sodium content of the experimentally reared butterflies with that of those collected on the wintering grounds in Mexico, as an estimate of what levels are typical of successfully migrating butterflies. Across male and females, there was no difference between our control and low-reared compared to butterflies collected from the wintering grounds; however, those from the high treatment did have significantly higher thoracic sodium than the wild-collected individuals (Figure 2). Together with the data on sodium excretion, these data suggest the high sodium levels are somewhat above the physiological optimum for monarchs, but are not exceptionally stressful.

We measured a range of other traits that might relate to migratory performance that similarly showed no variation across sodium treatments. We found that treatment did not influence wing coloration, including the proportion of black on the wing and orange coloration, though the orange color was influenced by developmental temperature (Figure S8), which has been found previously in monarchs [87]. Migratory monarchs generally have darker orange hue (redder) compared to non-migratory monarchs, and previous studies have found that within migratory monarchs, individuals with darker orange wings flew longer distances compared to individuals with lighter orange wings [63, 64, 81, 90]. Given that we found no difference in wing length or coloration, we would predict similar migratory flight performance across our different sodium treatments. Oe infections can also impact migratory success and survival, in part through changes to wing density and tensile strength which makes infected monarchs more prone to wing damage [67, 68]. We found no difference in Oe infections across our treatment groups, and generally detected a lower rate of Oe infection in our experimental butterflies than in the eastern North American monarch population (our study: 1.59%, other studies, <8% [91]) (Figure S9). It is possible that any potential negative impacts of sodium chloride were too conservative to detect with such low infection rates. Monarch migration and performance in flight experiments have also shown some more general connections to measures of immune investment. For example, monarchs with higher hemocyte counts have greater flight distances on flight mills [92], monarchs that experience diet restriction have decreases in both hemocyte concentrations and phenoloxidase activity, and increased phenoloxidase activity was found to correlate with adult lifespan [93]. Generally, immune measures in monarchs are decreased during reproduction compared to migration, but are generally robust during migration, even with high flight-related energy costs [94]. We found no variation in immune measures across our different sodium treatments (Figure S10), suggesting that sodium chloride was not impacting immune investment. We did detect some sex-specific variation in immune measures, which has also been found in other studies of monarchs (), where it has been suggested that there are different costs of investment in immune defenses for male and female monarch butterflies.

Overall, the findings regarding sodium are good news with respect to roadsides as potential habitat for monarchs (see also [54, 75]). Common milkweeds collected along most Minnesota roadsides generally have lower sodium values than the treatment groups here. Indeed, our low and high sodium chloride treatments correspond to 500 and 2000 ppm sodium, which is approximately the 90th and 99th percentiles of milkweeds collected from across Minnesota roadsides [33, 71]. It is important to note that the majority of roadside habitat in Minnesota falls along roads with fewer than 10,000 cars daily [95]. Milkweeds with levels 2000 ppm and higher are generally only seen on very highly trafficked roads (>30K per day) where salt application rates are higher, and close to the edge of the road [33]. Extrapolating these results to other locations will depend on road salt application rates, which vary widely across states, although are generally lower than in Minnesota [27]. In addition, recent efforts to reduce sodium chloride application rates will likely further reduce the chance of host plants with toxic levels of sodium. While our study manipulated sodium through spraying of plants, the contaminant content of roadside plants is a function of both plant uptake and dust on the leaf surface [96, 97]. It is possible that sodium within the leaf tissue is more stressful than that on the leaf surface, but previous lab experiments that manipulated sodium uptake found only minor reductions in monarch survival and body size with high levels of salt [75]. While the present study focused on salt as a possible toxic risk of roadside plants, it is important to note that there are other risks in roadside habitat, including metal exposure [54] and collisions with vehicles [98]; levels of heavy metals in most roadside milkweeds appear in safe limits to monarchs [54, 99], but the overall risks of collisions are less clear. At least with respect to the toxic risks of roadside milkweeds, research to date validates the utility of the average Midwestern roadside as habitat for monarchs, justifying costs of planting native milkweed in these areas [21].

## Acknowledgments

Thanks to Judie Prayfrock, Darcey Gans, Laura Wagner, Samantha Waddell, and Brandon Semke for help with rearing and butterfly processing. Thanks to Courtney Walker, Ashley Darst, Lexi Struble, Jeanna Edlund, and Brent Clanfield for help with specimens processing and filament images. Thanks to Dean Bower for hosting AKH in her lab to learn immune techniques.

## Data availability

All data and R code will be available on Mendeley.

## Supplemental Materials for

Road salt and monarch butterfly development: minimal effects on larval growth, immunity, wing coloration, and migration to Mexico.

## Supplemental Methods

### Rearing

We aimed to time our rearing with environmental conditions that would cue migration, but early enough to time release in late August, such that butterflies would move with the leading edge of migration. We first collected monarch eggs and larvae from the first wave of spring monarchs to move into Minnesota in early June, to form a group of breeder individuals for our experimental individuals. To raise the first generation, “breeding monarchs” monarch eggs, along with first and second instar larvae were collected from common milkweed on the University of Minnesota-St Paul campus, June 1-June 24 2019 and reared in 10 mesh “bug dorm” cages (61×61×61 cm) in ambient light in a greenhouse. Larvae were fed *ad lib* full-stalk cuttings of common milkweed, placed in water tubes (refilled daily if needed) and held upright in a plastic test tube rack. All milkweed was collected from weedy ditches, fields, and gardens across the St Paul campus.

To collect eggs for the experimental population (F2), breeder monarchs (the F1) were housed in large 24”x24”x36” mating cages (N=4-6 cages at any given time), with approximately 6 females and 4 males, with individuals replaced several times a week as they died. Butterflies were allowed to mature and mate for one week prior to the removal of all but one male and the addition of three milkweed stalks for egg laying. Butterflies had ad lib access to 10% honey water solution, available on a yellow sponge in a petri dish, replaced daily. Parental individuals were also screened for Oe.

We aimed to stagger egg-collecting and rearing to distribute the labor of feeding larvae and processing adult emergences. All experimental eggs (F2) were oviposited by breeder females (F1) onto stalks of common milkweed collected from Cedar Creek Research Station, held in water tubes in clear plastic bags at 4C (for up to two weeks) until use. Eggs collected from breeder females (and the resulting 1^st^ instar larvae) were housed in mesh cages, in a windowsill where they could be exposed to natural light (and natural declines in photoperiod). To keep milkweed plants from drying out, they were misted each day with distilled water and their water tubes were refilled. As larvae hatched, fresh milkweed was placed in the cages daily.

Larvae from a given egg collection cage were divided between the three salt treatments. Outdoors, individuals were reared to adulthood in mesh “bug-dorm” cages (61×61×61cm), fixed to the ground with tent stakes and string. Cages were placed under trees in areas that were mostly shaded to avoid overheating in direct sunlight (see Supplementary Figure 2). We aimed to transfer around 50-70 individual second instar larvae per cage. Larvae were transferred by moving individual milkweed plants or leaves into the new cages and counting all individuals on the plant. As discussed below, due to variation in counting accuracy of small larvae across workers varying in rearing experience, this method was less accurate than we had hoped.

To apply the salt treatment to a milkweed plant (prior to feeding to larvae), a stalk of milkweed (in a water tube) was sprayed with one squirt on the top and bottom of each leaf and left to dry for at least 15 minutes (outdoors) before being placed in larval rearing cages. We used 32-fl-oz (946 mL) Rubbermaid heavy duty spray bottles (3.3 ml/spray) to standardize spray flow rates. To avoid contamination from sweat, treatments were applied with gloved hands. Larvae in outdoor cages were fed at least once daily (generally mid-day depending on the weather). Cages with younger stage monarchs received 1-2 fresh stalks of milkweed daily, while cages with older stages received 4-10 milkweed plants daily, depending on their current consumption rate. We communicated with adjacent agricultural experiments to avoid pesticide application during monarch rearing. All milkweed used in rearing was collected from the Cedar Creek LTER field station. This site has consistently sandy soil, and, on average, low sodium concentrations in milkweeds (based on [1, 2]). In total, given rates of milkweed drying in our cages and consumption by larvae, we collected around 8000 stems of milkweed to rear our target 3000 butterflies. Harvested milkweed was immediately placed in water wicks containing distilled water (1-3 stems per tube) and stored in plastic bags at 4C for up to two weeks before use (see Supplementary Figure 2).

Adult butterflies emerging from outdoor cages were marked with a sharpie for processing on their first day, prior to tagging and release on their second number. Individual numbers were written 3-4 times on the wings to ensure the four-digit numbers could be read at a future date. On days with many emerging butterflies, butterflies were held between cage-removal and processing in glassine envelops stored in a dark cooler with ice packs. If butterflies emerged with deformed wings or otherwise could not fly at emergence (about 2.5% of individuals), they were sacrificed at –20C on the day of emergence, but were still tested for Oe. On an individuals’ release day, they received a Monarch-Watch tag, and their corresponding experimental number was recorded. Tags were color coded by treatment in the event that individual monarchs were sighted but not captured (although, in the end, all of our tag recoveries included full tag data). We colored tags with fabric markers (“Crafts for All” brand with Permanent German Ink); in a pilot comparison of four marker types on old tags, this brand did not fade in two months left out in the sun. At the end of the experiment, we carefully cleaned cages and quantified obviously diseased larva and pupa by counting any remaining carcasses.

To quantify wing length and color, we took standardized photos of butterflies using a lightbox (Supplementary Figure 4). To measure wing length, we used a standard measure that uses white dots on the thorax as landmarks for the base of the measurement, extending a line to the wing apex. Five people measured wing length for all butterflies without deformed wings; each person measured at least 150 butterflies and “measurer” was included as a covariate in analyses.

**Supplemental Figure 1.**
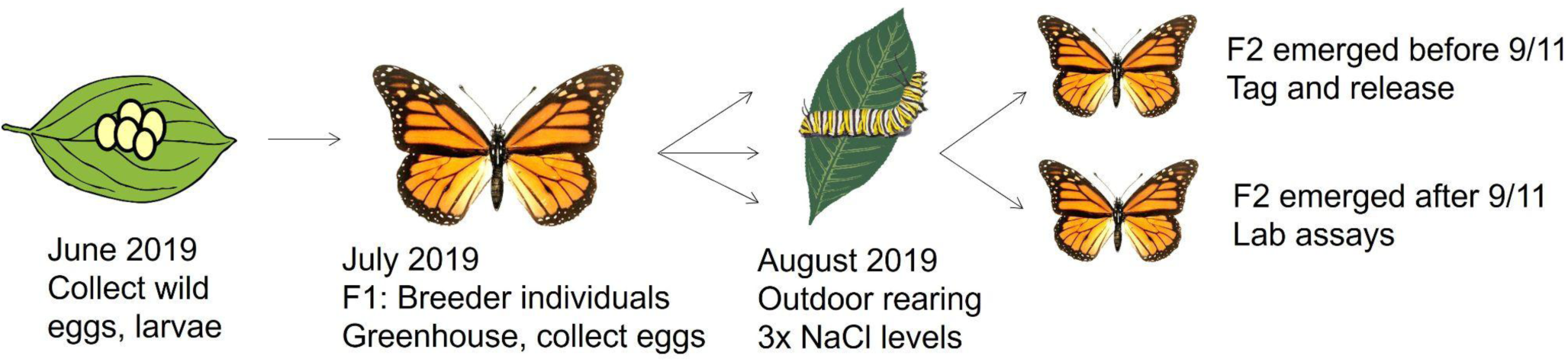
Overview of Rearing methods. We first reared a generation of breeder monarchs from wild collected eggs and larva. These individuals were reared in the greenhouse and served as breeding individuals to establish our experimental population. Larva from the second generation were divided between three salt treatments in outdoor cages. Individuals that emerged before 9/11 were tagged and released for migration. Individuals that emerged after 9/12 were used for various lab assays.

**Supplementary Figure 2.**
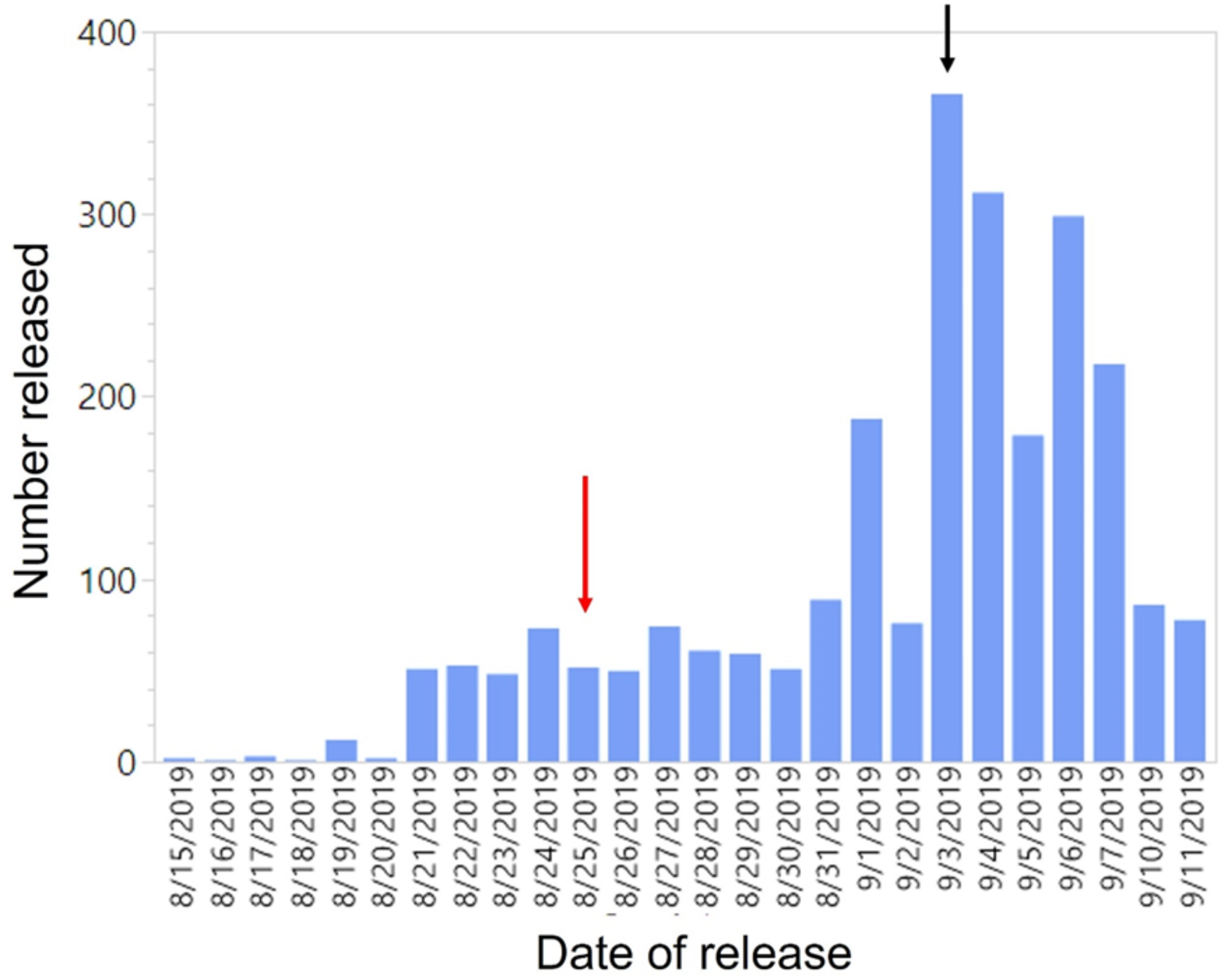
Summary of butterflies released. In our outdoor rearing, we were aiming for a peak release date around August 25^th^ (red arrow), when the leading edge of migration usually passes through the Twin Cities. However, due to cooler than average temperatures during the fall of 2019, our peak release date was delayed by nine days (black arrow). We reared a total of 2,959 butterflies and released a total of 2,464 tagged monarchs for migration.

**Supplementary Figure 3.**
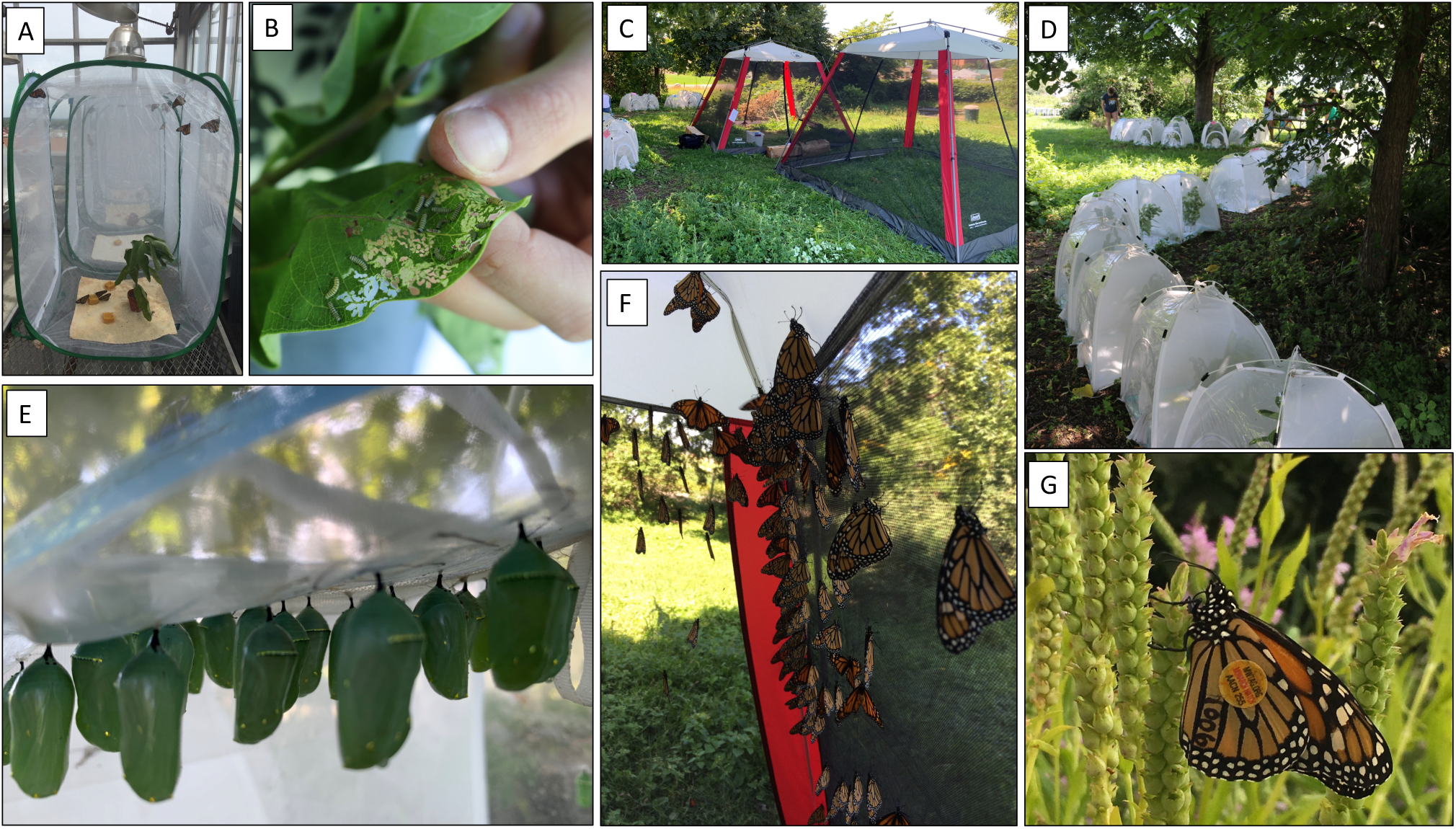
Rearing setup. Images of outdoor rearing protocol. Eggs from common-garden reared individuals were first collected on milkweed in the greenhouse (A). Early instar larvae (B) were transferred (on their host plants) to outdoor rearing cages (D) where they were fed ad lib until pupation (E). Emerged adults were given 24 hours in large flight cages for their wings to harden (C, F) before being tagged and released (G). Experimental butterflies were reared on common milkweed collected from one site at Cedar Creek Experimental Station. Milkweed stems were treated with either control, low, or high salt solutions and allowed to dry on test tube racks prior to placement in experimental rearing cages.

### Sodium analysis of butterfly tissue

We measured thoracic sodium through ICP-MS services at the Quantitative Bio-element Imaging Center at Northwestern University – the following is a shortened version of their protocol. Thorax samples were digested in 250 uL concentrated trace nitric acid (> 69%, Thermo Fisher Scientific, Waltham, MA, USA) and 62.5 uL concentrated trace hydrogen peroxide (>30.%, GFS Chemicals, Columbus, OH, USA) and placed at 65 °C for at least 3 hours to allow for complete sample digestion. Ultra-pure H2O (18.2 MΩ·cm) was then added to produce a final solution of 5.0% nitric acid (v/v) in a total sample volume of 5 mL. Quantitative standards for all elements were made using a 10 µg/mL mixed element standard (IV-ICPMS-71A, Inorganic Ventures, Christiansburg, VA, USA) which was used to create a 100 ng/mL mixed element standard in 5.0% nitric acid (v/v) in a total sample volume of 50 mL. An additional standard was prepared containing Na, Mg, P, Fe using single element standards (Inorganic Ventures, Christiansburg, VA, USA) which were combined to create a 5000 ng/mL mixed element standard in 5.0% nitric acid (v/v) in a total sample volume of 10 mL.ICP-MS was performed on a computer-controlled (QTEGRA software) Thermo iCapQ ICP-MS (Thermo Fisher Scientific, Waltham, MA, USA) operating in KED mode and equipped with an ESI SC-2DX PrepFAST autosampler (Omaha, NE, USA). Internal standard was added inline using the prepFAST system and consisted of 1 ng/mL of a mixed element solution containing Bi, In, 6Li, Sc, Tb, Y (IV-ICPMS-71D from Inorganic Ventures). Online dilution was also carried out by the prepFAST system and used to generate calibration curves consisting of 100, 50, 10, 5, 1 and 0.5 ng/g of all elements, and 5000, 2500, and 1000 ng/g additional calibration points for Na, Mg, P, and Fe. Each sample was acquired using 1 survey run (10 sweeps) and 3 main (peak jumping) runs (40 sweeps). The isotopes selected for analysis were 23Na, 24Mg, 27Al, 31P, 39K, 56Fe, 57Fe, 60Ni, 63Cu, 65Cu, 66Zn, 111Cd, and 208Pb and 89Y, 115In (chosen as internal standards for data interpolation and machine stability). Instrument performance is optimized daily through autotuning followed by verification via a performance report.

### Weather Data

Vapor pressure deficit (VPD, measured in pounds per square inch, psi) accounts for both temperature and humidity as it is a measure of the amount of water vapor in the air (actual vapor pressure) compared to how much capacity there is for water vapor at a given temperature (saturation vapor pressure, SVP). Where *T*= temperature and *RH =* relative humidity, we calculated VPD as:

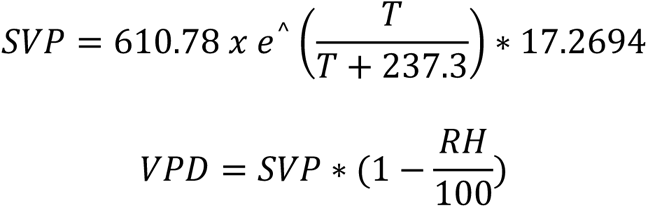

In our dataset, VPD was correlated with both temperature and humidity (Supplementary Figure 4 AB), thus we only looked at VPD in our models. Both VPD and temperature were also both correlated with emergence date, which we included in our models as a random effect (Supplementary Figure 4 CD).

**Supplementary Figure 4.**
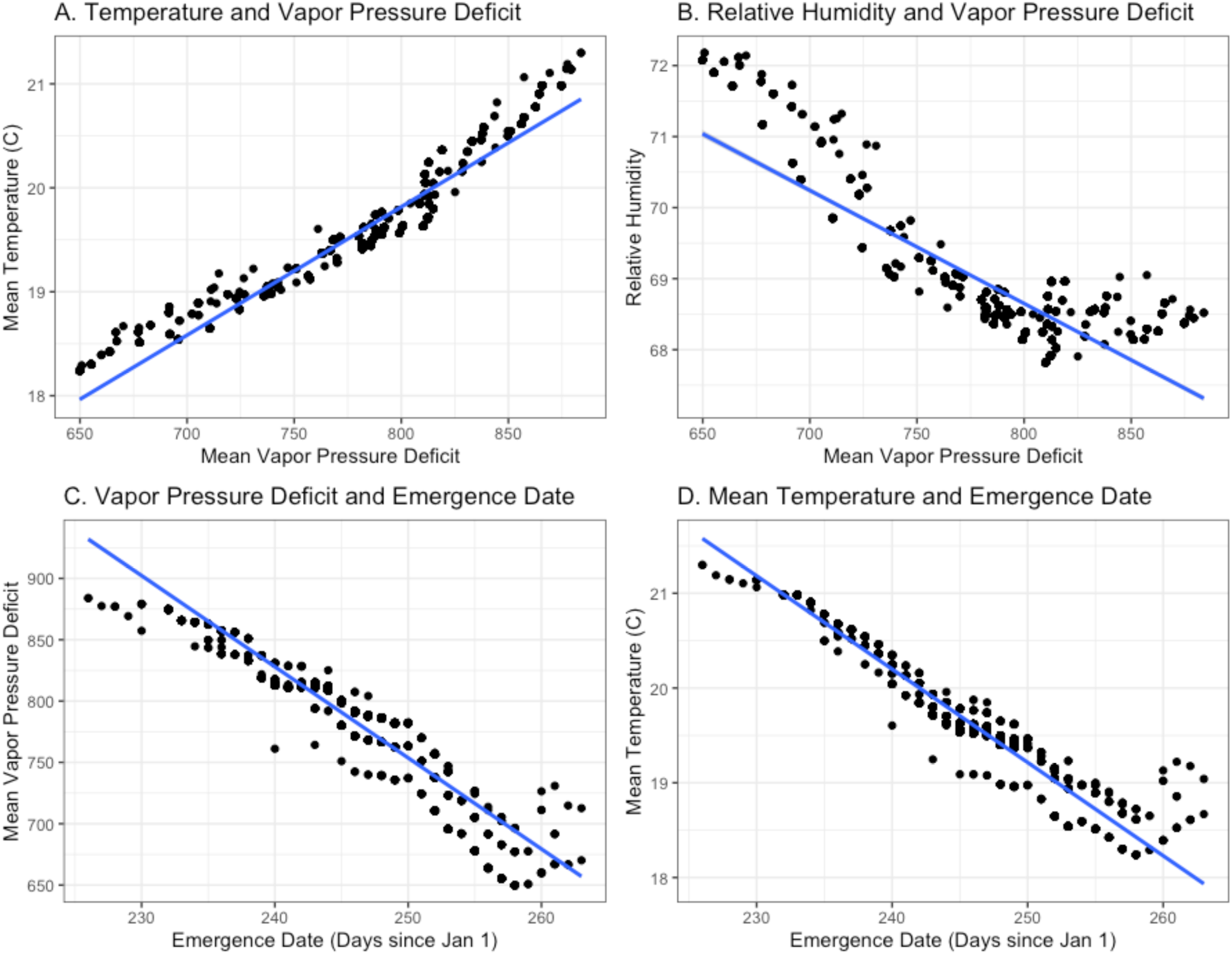
Vapor Pressure Deficit. (A) Temperature and VPD were tightly correlated in our dataset, with higher VPD at warmer temperatures. (B) Relative Humidity and VPD were correlated, with higher VPD being drier. (C) VPD was also correlated with emergence data, where earlier dates were warmer and drier compared to later dates, which were cooler and wetter. (D) Temperature was correlated with emergence data, where earlier dates were warmer than later dates. Emergence dates range from 8/14/19 (Julian date 226) to 9/20/19 (Julian date 263).

### Morphological measurements and wing coloration

We measured wing length using a standard wing length measurement used by Monarch Watch to calibrate our measurements with other studies (Supplementary Figure 4C). For wing coloration measures, we extracted three distinct color variables: proportion of black, hue of orange and saturation of orange. To estimate proportion of black on the wings, we used the color adjacency function in pavo2.0 package in R. This function allows us to subtract the background, crop a Region of Interest (ROI, Supplementary Figure 4D) and classify all pixels within the ROI into three distinct categories: black, orange, and white, with references to black, orange, and white color standards using K-means clustering. The resulting output was then used to calculate the proportion of black on wings using the following formula: # black pixels / (# black pixels + # orange pixels + # white pixels). To estimate hue and saturation of orange on the wings, we extracted average Red (R), Green (G), and Blue (B) values for orange regions within the RIO in MATLAB (R2022). Subsequently, R, G, B values were averaged for every individual and hue and saturation were calculated. Since all individuals were photographed under controlled light conditions in a light box, no additional standardization was performed.

**Supplementary Figure 5.**
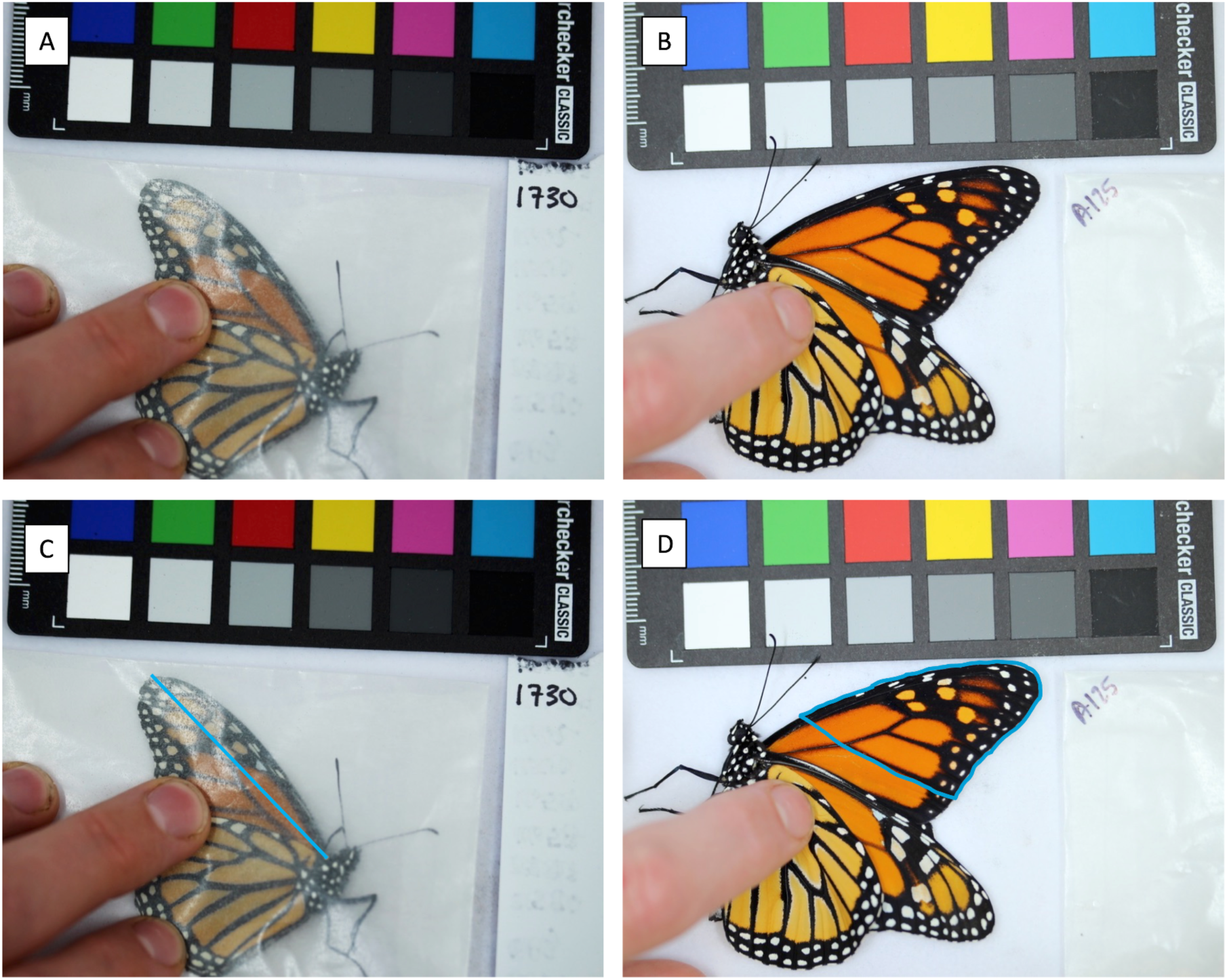
Wing photos. (A) Standardized wing photo that we took to measure wing length, in a glassine envelope to hold wings flat for accurate measurement. (B) Standardized wing photo that we took to measure wing color of dorsal forewing. (C) Landmarks that we used to measure wing length as a proxy for body size. (D) Landmarks that we used to quantify wing coloration: proportion of black on the wing and the saturation and hue of the orange coloration in wing cells.

### Immune measurements

Individuals were held for <u>48 hrs</u> after eclos<u>ion</u> to allow their immune system to mature. During these two days, butterflies were kept in cages indoors near a window with *ad lib* access to 10% honey water. We chilled butterflies on ice for 1-2 minutes and, under a dissecting microscope, made a small puncture using a sterilized insect pin on the side of the abdomen. We then collected hemolymph using 5ul capillary tubes. 5ul of hemolymph was immediately added to 20ul of anticoagulant buffer (10mM EDTA, 10mM citric acid, in PBS, pH 7.4) and placed on ice for hemocyte counts. The other 5ul of hemolymph was added to 250 ul of 10mM chilled sodium cacodylate buffer (NaCac) and stored at –80C for phenoloxidase assays. We then inserted a 2mm knotted <u>nylon</u> filament made with roughened, sterilized, 0.2mm diameter clear monofilament fishing line (<u>Berkley ®</u> Trilene) into the wound by sliding it laterally just under the cuticle. The knot held the inserted filament in place and allowed for easy retrieval. To allow butterflies to react to the filament, we placed them in glassine envelopes and held them in a 24C incubator for 1 hour. Previous pilot studies indicated that 1 hour allowed for a detectable immune response and best captured variation between individuals (longer time periods allowed most individuals to completely melanize the filament). After retrieval, filaments were stored in dry tubes at –80 and butterflies were frozen.

Inserted filaments simulate an immune challenge and allow us to measure the encapsulation response. During the encapsulation process, hemocytes aggregate around the foreign object, creating a darkened melanized capsule. This process can kill invaders that are too large to phagocytose (i.e. parasitoid eggs, fungal pathogens) through oxidative damage, starvation, or cytotoxic molecules [3]. To quantify the encapsulation response, we photographed filaments with a black and white color standard using Leica Stereo microscope at 20× magnification and used Adobe photoshop to quantify the degree of melanization (darkness of the filament) following [4] (see supplement figure 5).

We also quantified the density of hemocytes in the hemolymph using a Bright-Line^TM^ hemocytometer and a light microscope at 40× magnification. Hemocytes are a central component of the insect immune system and are involved in both humoral and cellular immune responses, including phagocytosis and encapsulation. Higher hemocyte density has been associated with increased immune capacity and parasite resistance [5]. Finally, we measured phenoloxidase (PO) and prophenoloxidase (proPO) enzymatic activity and quantified hemolymph protein. PO is central to many functions of the insect immune response. It initiates the melanization response, which is important for encapsulation, plays a role in coordinating the cellular response, and is involved in humoral immunity and antimicrobial activity in the hemolymph. PO levels have been connected to variation in parasite resistance in a number of species [6]. PO comes in both an activated form and an inactivated or bound form, ProPO. The pool of bound ProPO can be quickly activated during an immune response. To quantify ProPO, we used chymotrypsin (20mg/ml) to convert ProPO to PO (we could then subtract our measure of free PO from total PO to estimate ProPO). We determined the activity of PO using an absorbance spectrometer plate reader (Tecan) at 490nm using 2mM dopamine as a substrate. We measured the level of protein in the hemolymph samples as a way to further standardize our PO measures using a Bradford Coomassie Plus kit (Fisher Scientific). We then calculated the activity of PO per min per ug of protein as a measure of immune capacity.

**Supplementary Figure 6.**
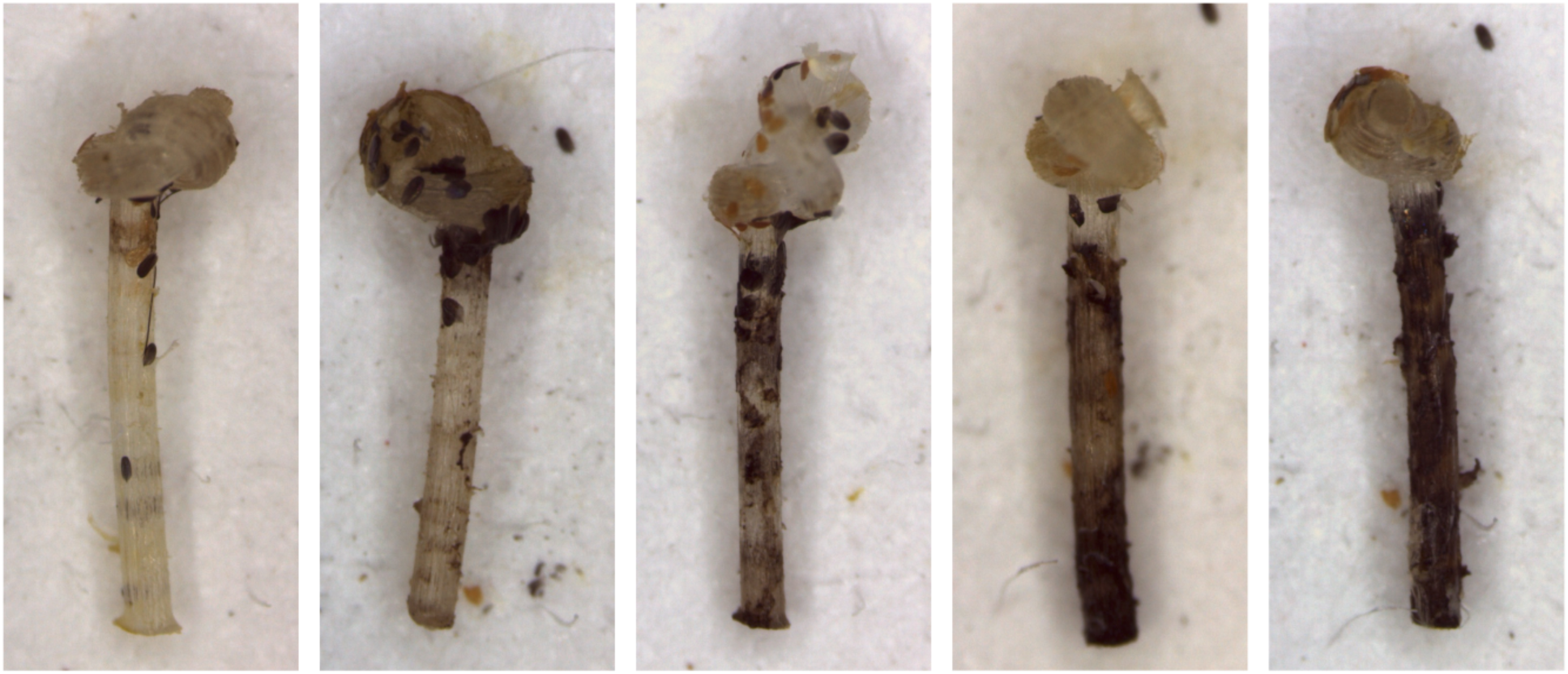
Melanized filaments. Examples of variation in the degree of encapsulation response (melanization) to inserted filaments as a measure of immune capacity from weak (left) to strong (right). We quantify the degree of melanization (darkness of the filament) relative to a black and white standard in each photograph (not visible in the cropped images above) using Adobe photoshop.

## Supplemental Results

### Growth and development

**Table S1.** Results from linear mixed-effects model for wing length including both sexes. *N=2,807 adult monarchs*.

|  |  |  |  |  |
| --- | --- | --- | --- | --- |
| *Term* | *Coefficient* | *SE* | *t-value* | *p* |
| Treatment (H) | -0.42 | 0.32 | -1.31 | 0.19 |
| Treatment (L) | 0.29 | 0.31 | 0.93 | 0.35 |
| Mean VPD | -0.01 | 0.002 | -4.04 | < 0.001 |
| Sex (M) | 0.64 | 0.09 | 6.94 | < 0.001 |

**Table S2.** Results from linear mixed-effects model for wing length for females. *N=1,458 adult monarchs*.

| *Term* | *Coefficient* | *SE* | *t-value* | *p* |
| --- | --- | --- | --- | --- |
| Treatment (H) | -0.36 | 0.31 | -1.14 | 0.25 |
| Treatment (L) | -0.08 | 0.31 | -0.27 | 0.79 |
| Mean VPD | -0.01 | 0.002 | -3.51 | < 0.001 |

**Table S3.** Results from linear mixed-effects model for wing length for males. *N=1,349 adult monarchs*.

| *Term* | *Coefficient* | *SE* | *t-value* | *p* |
| --- | --- | --- | --- | --- |
| Treatment (H) | -0.42 | 0.32 | -1.31 | 0.19 |
| Treatment (L) | 0.29 | 0.31 | 0.93 | 0.35 |
| Mean VPD | -0.01 | 0.002 | -4.04 | < 0.001 |

**Table S4.** Results from linear mixed-effects model for development time (days from egg to eclose). *N=2,954 adult monarchs*.

| *Term* | *Coefficient* | *SE* | *t-value* | *p* |
| --- | --- | --- | --- | --- |
| Sex (M) | -0.002 | 0.004 | -0.36 | 0.72 |
| Treatment (H) | -0.004 | 0.11 | -0.04 | 0.97 |
| Treatment (L) | -0.01 | 0.15 | -0.07 | 0.94 |
| Mean VPD | -0.152 | 0.002 | -65.40 | < 0.001 |

**Table S5.**
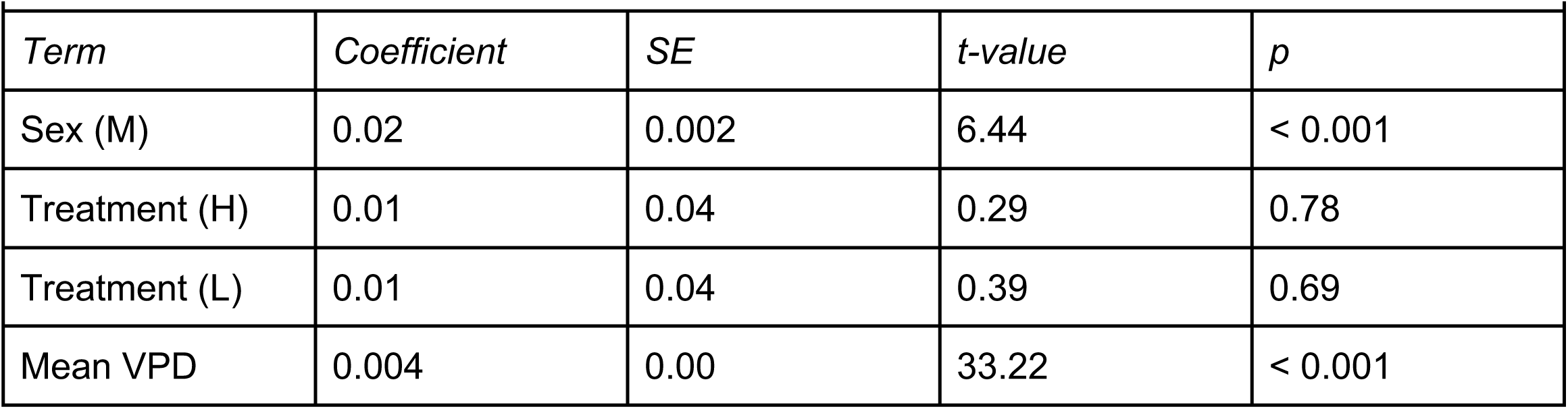
Results from linear mixed-effects model for growth rate (wing length / development time) including both sexes. *N=2,807 adult monarchs*.

| *Term* | *Coefficient* | *SE* | *t-value* | *p* |
| --- | --- | --- | --- | --- |
| Sex (M) | 0.02 | 0.002 | 6.44 | < 0.001 |
| Treatment (H) | 0.01 | 0.04 | 0.29 | 0.78 |
| Treatment (L) | 0.01 | 0.04 | 0.39 | 0.69 |
| Mean VPD | 0.004 | 0.00 | 33.22 | < 0.001 |

**Table S6.** Results from linear mixed-effects model for growth rate (wing length / development time) for females. *N= 1,458 adult monarchs*.

| *Term* | *Coefficient* | *SE* | *t-value* | *p* |
| --- | --- | --- | --- | --- |
| Treatment (H) | 0.02 | 0.04 | 0.58 | 0.56 |
| Treatment (L) | 0.01 | 0.04 | 0.33 | 0.74 |
| Mean VPD | 0.004 | 0.00 | 27.56 | < 0.001 |

**Table S7.**
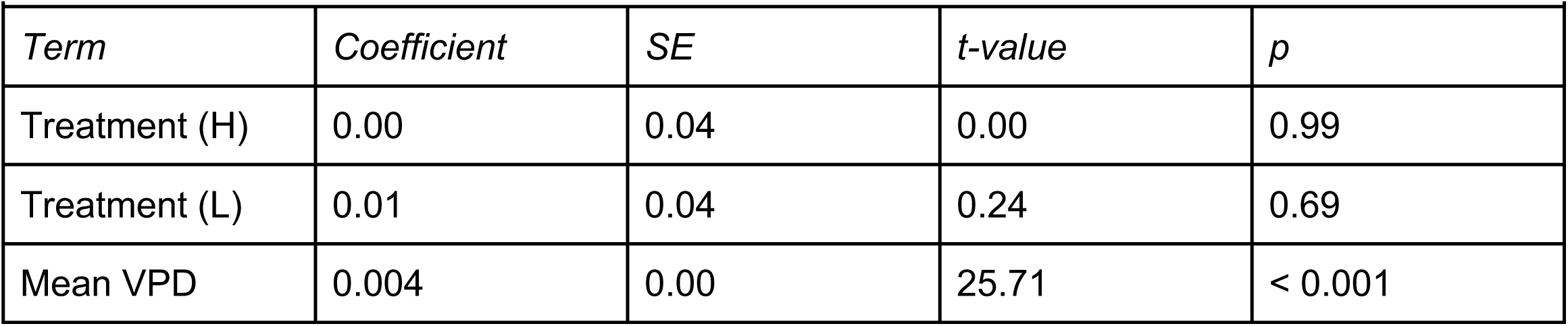
Results from linear mixed-effects model for growth rate (wing length / development time) males. *N= 1,349 adult monarchs*.

**Supplementary Figure 7.**
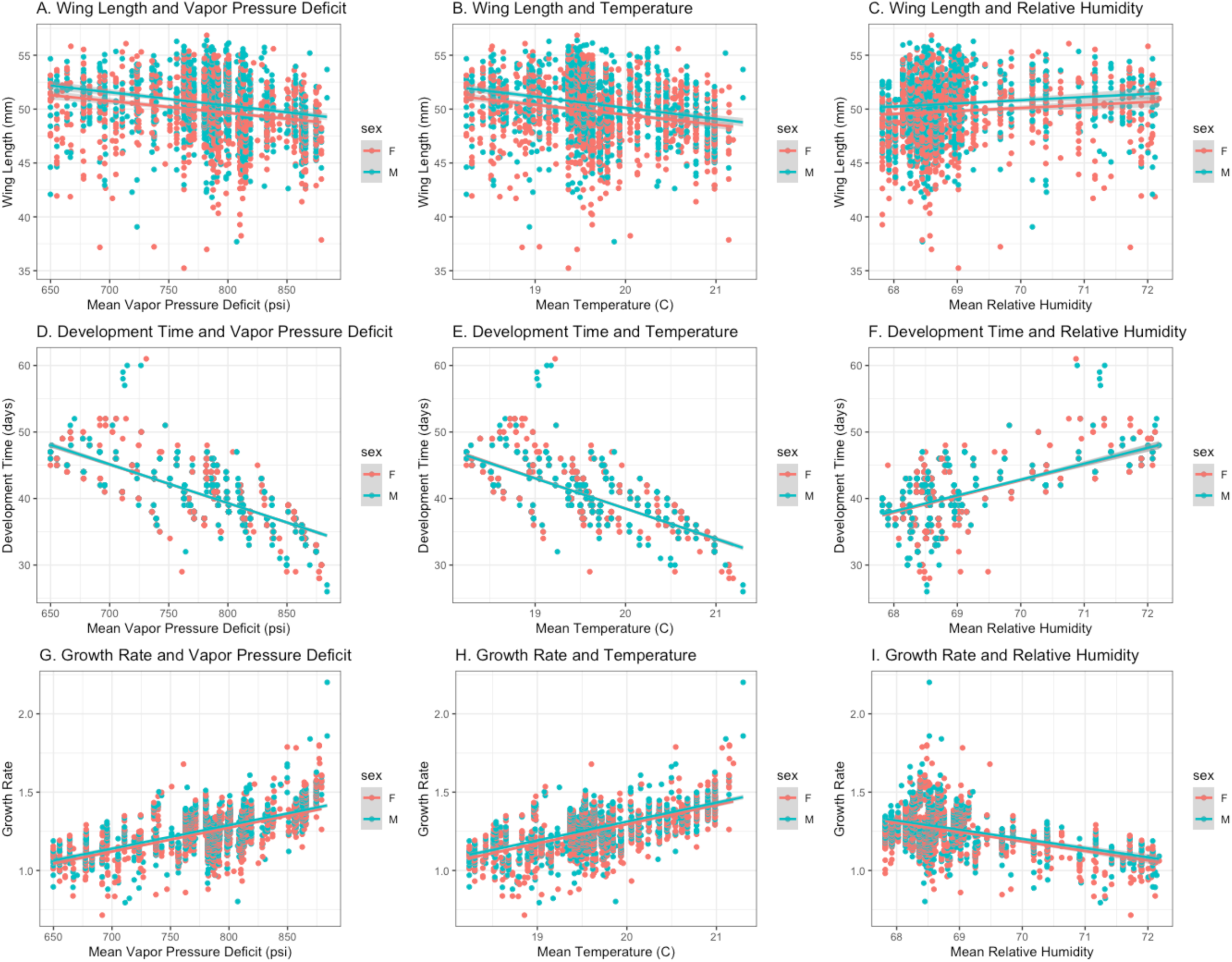
Body size and development time. Wing length decreased as mean vapor pressure deficit (VPD) increased (A), corresponding to warmer temperatures (B) and less humidity (C). Males (blue) had longer wings, on average, compared to females (red). Development time (number of eggs from egg to eclose) decreased as mean VPD increased (D), corresponding to warmer temperature (E) and less humidity (F) There was no difference in development time between males and females. Growth rate (wing length / development time) increased as mean VPD increased (G), corresponding to warmer temperatures (H) and more humid conditions (I). Males had faster growth rates compared to females.

### Survival in the field and lab

**Table S8.**
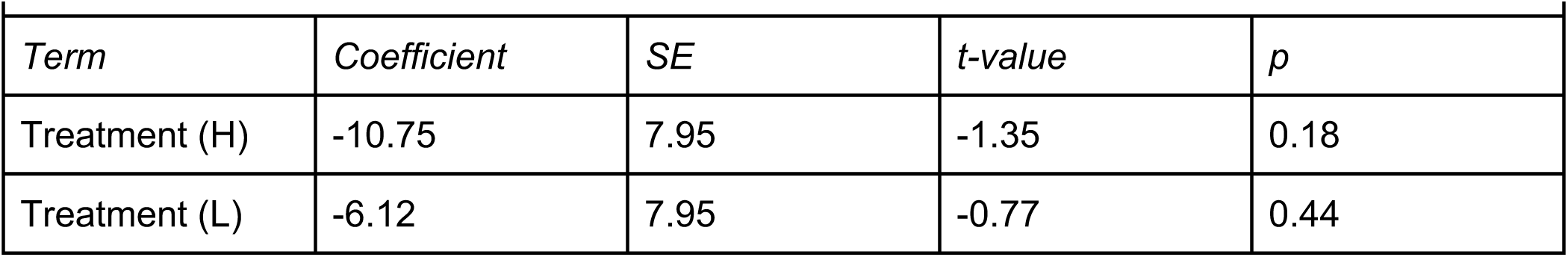
Results from mixed-effects model for outdoor survival (cage level). *N= 60 cages*.

**Table S9.** Results from mixed-effects model for proportion of diseased larva (cage level). *N= 60 cages*.

| *Term* | *Coefficient* | *SE* | *z-value* | *p* |
| --- | --- | --- | --- | --- |
| Treatment (H) | -0.33 | 0.27 | -1.23 | 0.22 |
| Treatment (L) | -0.48 | 0.27 | -1.81 | 0.07 |

**Table S10.**
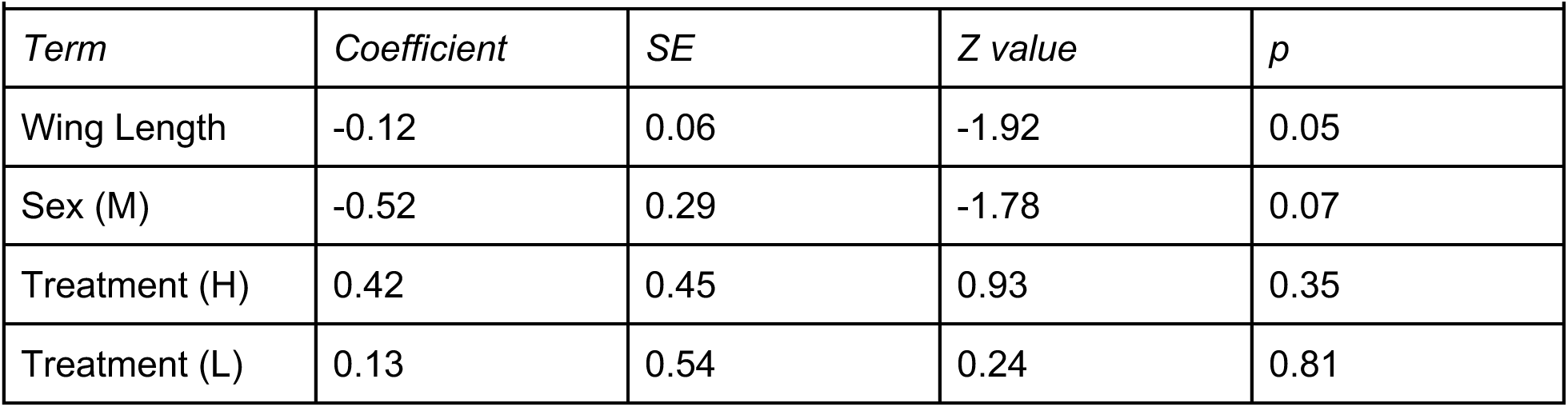
Results from cox mixed-effects model for indoor survival in the constant climate chamber (21C). *N= 60 butterflies*.

**Supplementary Figure 8.**
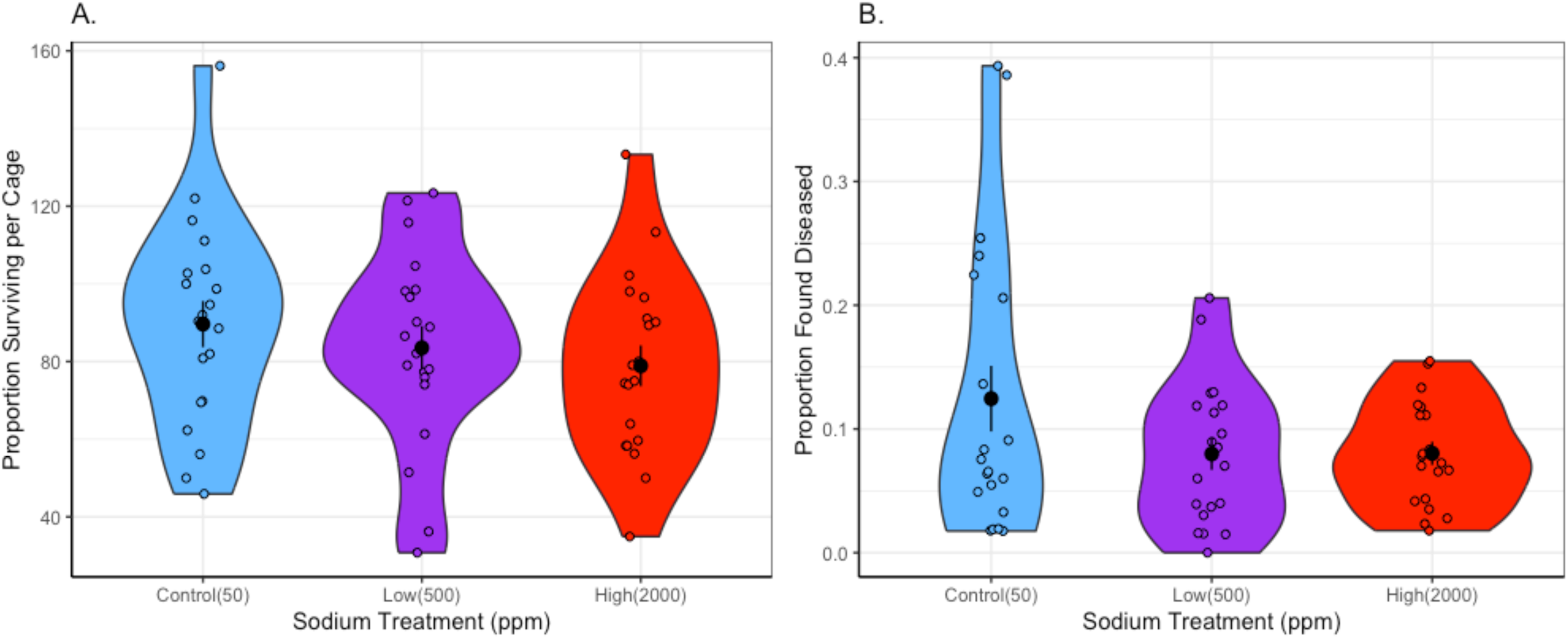
Outdoor survival. (A) proportion of larva that survived to adult butterflies for each treatment in our outdoor cages. (B) Proportion of larva found to be obviously diseased in cages at the end of the experiment for each treatment group. We note that these data are quite noisy as counts of larva being put into outdoor cages was not very accurate.

**Table S11.** Results from cox mixed-effects model for indoor survival in the variable climate chamber (21C, 7C). *N= 90 butterflies*.

| *Term* | *Coefficient* | *SE* | *Z value* | *p* |
| --- | --- | --- | --- | --- |
| Wing Length | -0.20 | 0.04 | -4.72 | < 0.001 |
| Sex (M) | 0.05 | 0.24 | 0.21 | 0.83 |
| Treatment (H) | 0.42 | 0.44 | 0.67 | 0.33 |
| Treatment (L) | 0.28 | 0.46 | 0.61 | 0.54 |

**Supplementary Figure 9.**
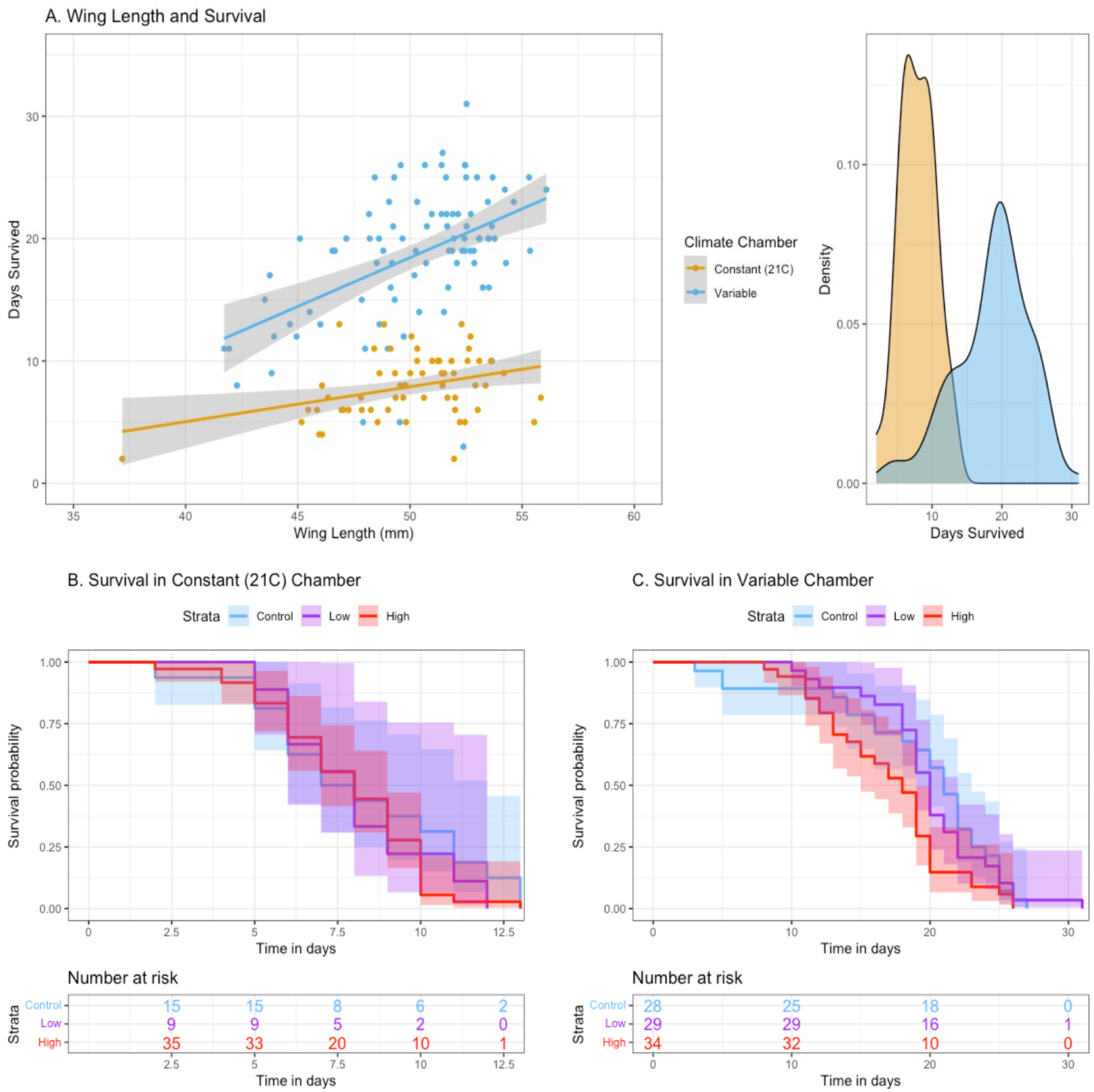
Survival in the lab. (A) Butterflies with larger body sizes (longer wing length) survived better in both the variable (21C, 7C) and constant (21C) climate chambers. As seen in the density plot, butterflies in the variable climate chamber lived longer compared to butterflies in the constant climate chamber (constant: mean = 7.92, SD = 2.56; variable: mean = 18.63, SD = 5.34). We found no significant effect of treatment on survival in either the constant chamber (B) or the variable chamber (C).

### Wing Coloration

**Table S12.** Results from a mixed-effects model for the proportion of black on the wing for females. *N= 179 butterflies*.

|  |  |  |  |  |
| --- | --- | --- | --- | --- |
| *Term* | *Coefficient* | *SE* | *Z value* | *p* |
| Treatment (H) | -0.01 | 0.02 | -0.34 | 0.73 |
| Treatment (L) | 0.01 | 0.02 | 0.22 | 0.79 |
| Mean Temp | 0.02 | 0.02 | 0.02 | 0.15 |

**Table S13.** Results from a mixed-effects model for the proportion of black on the wing for males. *N= 176 butterflies*.

|  |  |  |  |  |
| --- | --- | --- | --- | --- |
| *Term* | *Coefficient* | *SE* | *Z value* | *p* |
| Treatment (H) | 0.01 | 0.02 | -0.36 | 0.72 |
| Treatment (L) | -0.02 | 0.02 | -1.03 | 0.30 |
| Mean Temp | 0.04 | 0.03 | 1.68 | 0.09 |

**Table S14.** Results from a mixed-effects model for orange hue for females. *N= 179 butterflies*.

|  |  |  |  |  |
| --- | --- | --- | --- | --- |
| *Term* | *Coefficient* | *SE* | *T value* | *p* |
| Treatment (H) | -0.24 | 0.63 | -0.39 | 0.69 |
| Treatment (L) | -1.05 | 0.63 | -1.66 | 0.10 |
| Mean Temp | -1.45 | 0.42 | -3.43 | < 0.001 |

**Table S15.**
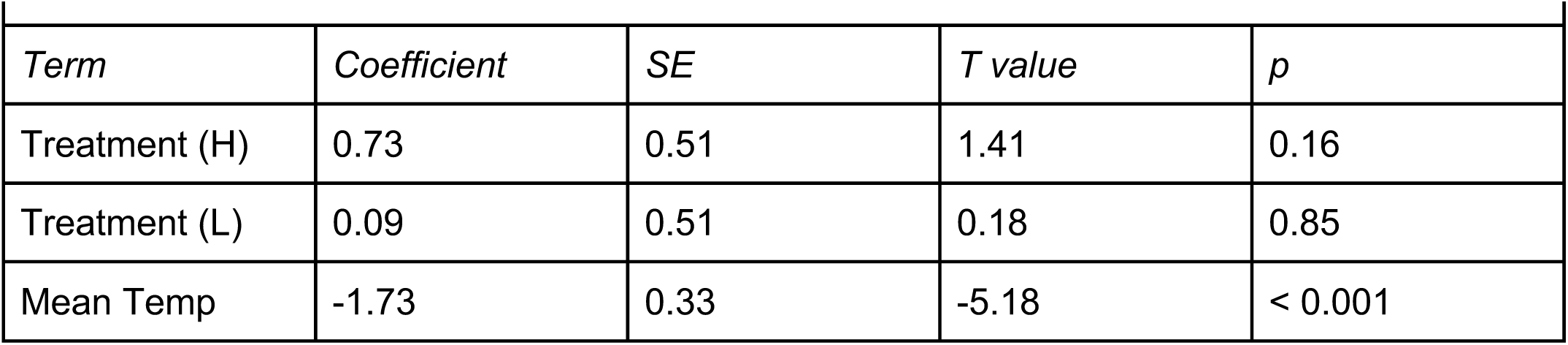
Results from a mixed-effects model for orange hue for males. *N= 176 butterflies*.

| *Term* | *Coefficient* | *SE* | *T value* | *p* |
| --- | --- | --- | --- | --- |
| Treatment (H) | 0.73 | 0.51 | 1.41 | 0.16 |
| Treatment (L) | 0.09 | 0.51 | 0.18 | 0.85 |
| Mean Temp | -1.73 | 0.33 | -5.18 | < 0.001 |

**Table S16.** Results from a mixed-effects model for orange saturation for females. *N= 179 butterflies*.

| *Term* | *Coefficient* | *SE* | *T value* | *p* |
| --- | --- | --- | --- | --- |
| Treatment (H) | -0.25 | 0.97 | -0.26 | 0.76 |
| Treatment (L) | 0.37 | 0.97 | 0.39 | 0.70 |
| Mean Temp | -2.70 | 0.65 | -4.16 | < 0.001 |

**Table S17.** Results from a mixed-effects model for orange saturation for males. *N= 176 butterflies*.

| *Term* | *Coefficient* | *SE* | *T value* | *p* |
| --- | --- | --- | --- | --- |
| Treatment (H) | -0.35 | 0.67 | -0.52 | 0.60 |
| Treatment (L) | -0.56 | 0.65 | -0.85 | 0.40 |
| Mean Temp | -0.83 | 0.43 | -1.93 | 0.055 |

**Supplementary Figure 10.**
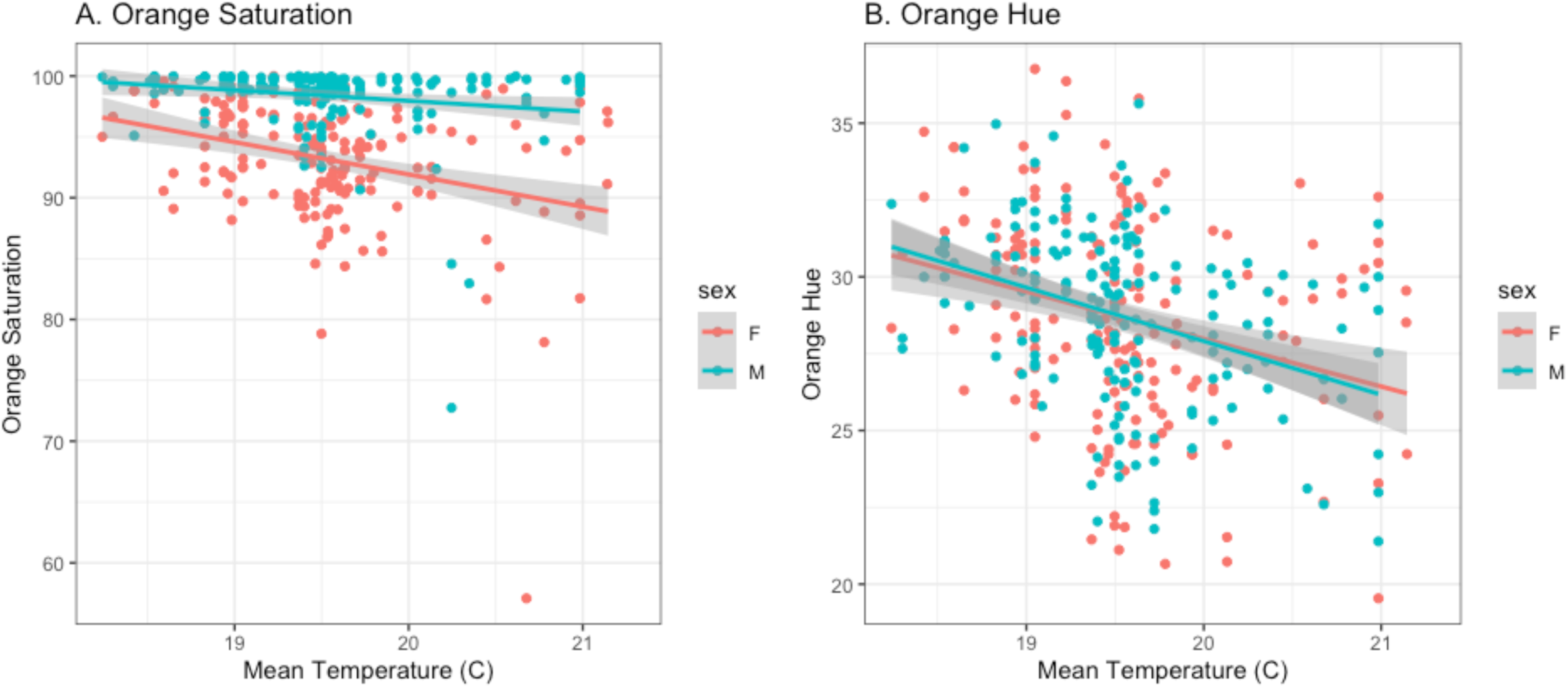
Orange wing coloration and temperature. (A) Orange saturation decreased for both males and females as mean temperature during development increased. (B) Orange hue decreased for both males and females as mean temperature during development increased.

### Ophryocystis elektroscirrha Infections

**Table S18.** Results from a mixed-effects model for OE infections (presence / absence). *N= 2,959 butterflies*.

|  |  |  |  |  |
| --- | --- | --- | --- | --- |
| *Term* | *Coefficient* | *SE* | *Z value* | *p* |
| Treatment (H) | -0.67 | 1.71 | -0.39 | 0.70 |
| Treatment (L) | 4.87 | 3.18 | 1.53 | 0.13 |
| Sex (M) | -0.39 | 0.58 | -0.68 | 0.50 |

**Supplementary Figure 11.**
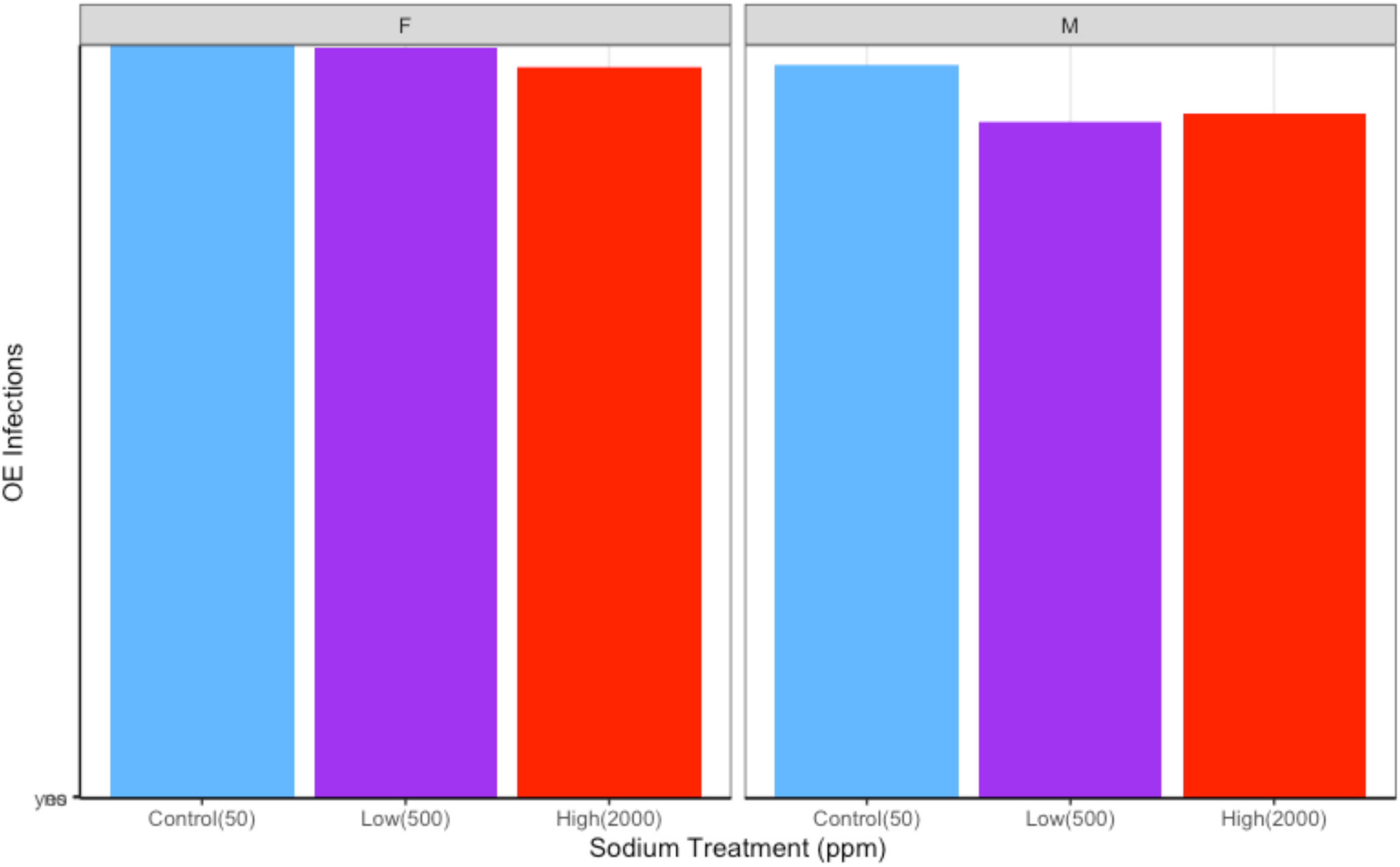
Ophryocystis elektroscirrha Infections. Presence / absence of OE infections across treatment groups for both females (first panel) and males (second panel). There was no effect of treatment on OE infections.

### Immune Measures

**Table S19.** Results from the full mixed-effects model for hemocyte counts (including sex:treatment interaction). *N= 75 butterflies*.

|  |  |  |  |  |
| --- | --- | --- | --- | --- |
| *Term* | *Coefficient* | *SE* | *Z value* | *p* |
| Sex (M) | 0.58 | 0.04 | -13.46 | < 0.001 |
| Treatment (H) | 0.49 | 0.26 | 1.85 | 0.06 |
| Treatment (L) | 0.04 | 0.25 | 0.18 | 0.86 |
| Sex(M) :<br>Treatment (H) | -0.89 | 0.05 | -17.46 | < 0.001 |
| Sex(M) :<br>Treatment (L) | -0.47 | 0.05 | -9.68 | < 0.001 |

**Table S20.** Results from a mixed-effects model for hemocyte counts for females. *N= 38 butterflies*.

| *Term* | *Coefficient* | *SE* | *Z value* | *p* |
| --- | --- | --- | --- | --- |
| Treatment (H) | 0.03 | 0.23 | 0.15 | 0.88 |
| Treatment (L) | 0.20 | 0.23 | 0.90 | 0.37 |

**Table S21.** Results from a mixed-effects model for hemocyte counts for males. *N= 37 butterflies*.

| *Term* | *Coefficient* | *SE* | *Z value* | *p* |
| --- | --- | --- | --- | --- |
| Treatment (H) | -0.25 | 0.50 | -0.50 | 0.62 |
| Treatment (L) | -0.38 | 0.47 | -0.81 | 0.42 |

**Table S22.** Results from the full mixed-effects model for filament encapsulation. *N= 66 butterflies*.

| *Term* | *Coefficient* | *SE* | *T value* | *p* |
| --- | --- | --- | --- | --- |
| Sex (M) | 1.74 e-05 | 1.87 e-04 | 0.09 | 0.93 |
| Treatment (H) | -8.15 e-05 | 2.60 e-04 | -0.31 | 0.75 |
| Treatment (L) | 1.39 e-04 | 2.44 e-04 | 0.57 | 0.57 |

**Table S23.** Results from the full mixed-effects model for PO. *N= 71 butterflies*.

| *Term* | *Coefficient* | *SE* | *T value* | *p* |
| --- | --- | --- | --- | --- |
| Sex (M) | 0.24 | 0.14 | 1.68 | 0.10 |
| Treatment (H) | -0.17 | 0.20 | -0.89 | 0.38 |
| Treatment (L) | -0.17 | 0.19 | -0.91 | 0.37 |

**Table S24.**
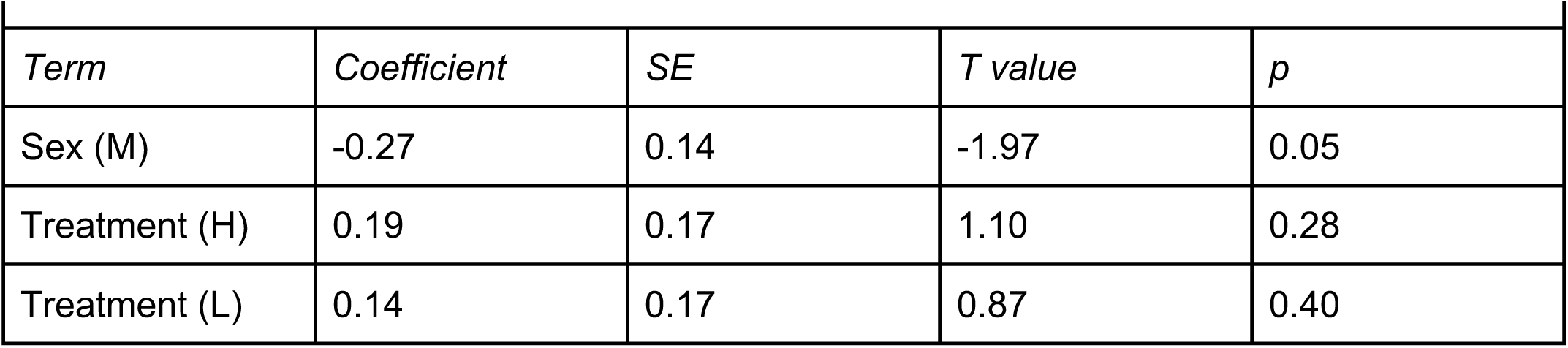
Results from the full mixed-effects model for ProPO. *N= 68 butterflies*.

| *Term* | *Coefficient* | *SE* | *T value* | *p* |
| --- | --- | --- | --- | --- |
| Sex (M) | -0.27 | 0.14 | -1.97 | 0.05 |
| Treatment (H) | 0.19 | 0.17 | 1.10 | 0.28 |
| Treatment (L) | 0.14 | 0.17 | 0.87 | 0.40 |

**Table S25.** Results from the full mixed-effects model for ProPO for females. *N= 36 butterflies*.

| *Term* | *Coefficient* | *SE* | *T value* | *p* |
| --- | --- | --- | --- | --- |
| Treatment (H) | 0.20 | 0.24 | 0.82 | 0.42 |
| Treatment (L) | 0.39 | 0.24 | 1.66 | 0.12 |

**Table S26.** Results from the full mixed-effects model for ProPO for males. *N= 32 butterflies*.

| *Term* | *Coefficient* | *SE* | *T value* | *p* |
| --- | --- | --- | --- | --- |
| Treatment (H) | 0.19 | 0.24 | 0.78 | 0.45 |
| Treatment (L) | -0.13 | 0.23 | -0.56 | 0.58 |

**Supplementary Figure 12.**
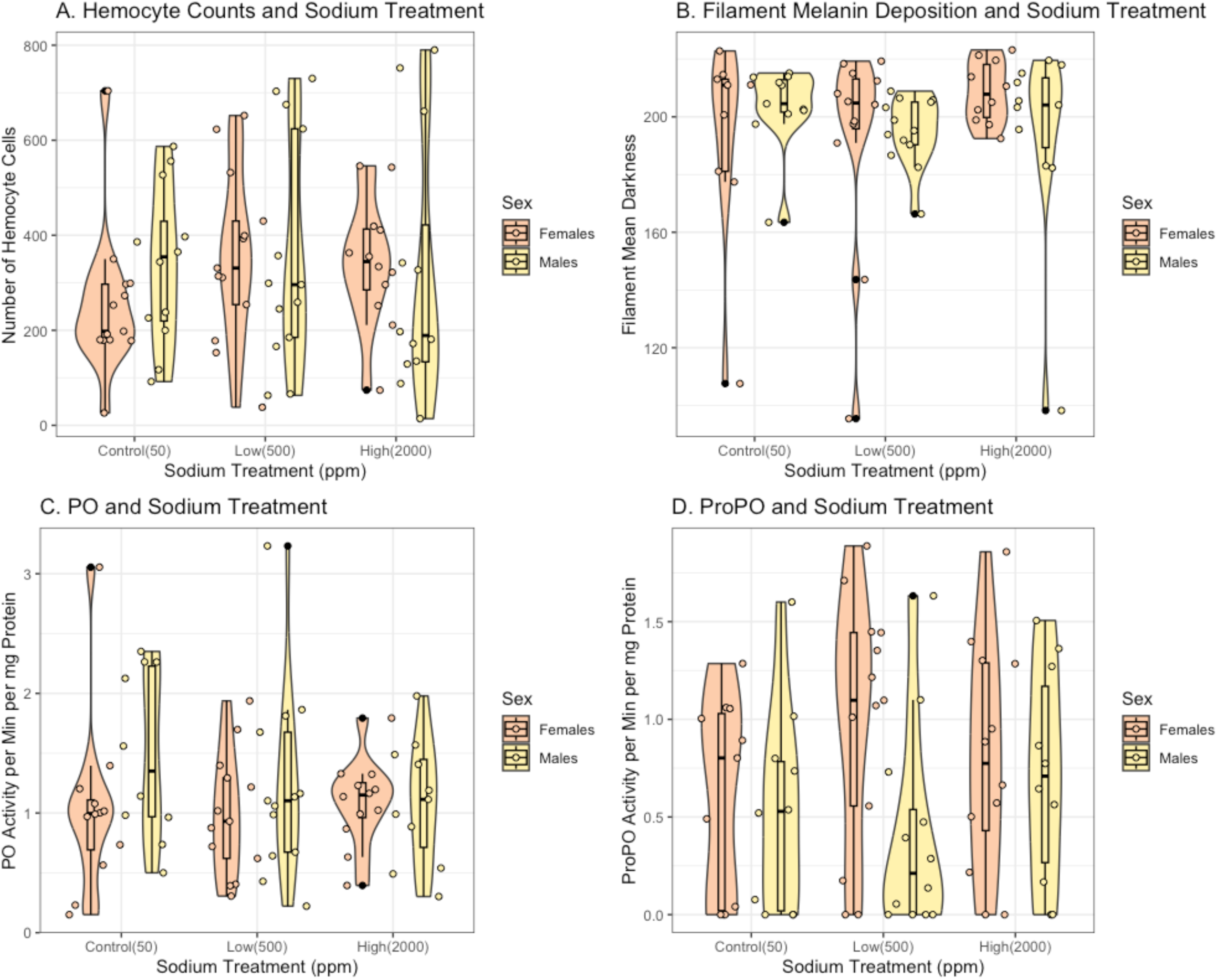
Immune Assays. Four standard immune assays separated by sex and treatment group. (A) Hemocyte counts, (B) Filament melanin deposition, (C) Phenoloxidase activity, and (D) ProPhenoloxidase activity. We found no significant effect of treatment for any of these immune measures.

